# Diphenyleneiodonium chloride inhibits MYCN-amplified neuroblastoma by targeting MYCN induced mitochondrial alterations

**DOI:** 10.1101/2024.10.20.619268

**Authors:** Soraya Epp, Stephanie Maher, Amirah Adlina Abdul Aziz, Simone Marcone, Donagh Egan, Saija Haapa-Paananen, Vidal Fey, Kristiina Iljin, Kieran Wynne, Lasse D. Jensen, Walter Kolch, Melinda Halasz

**Affiliations:** Systems Biology Ireland, School of Medicine, University College Dublin, Dublin, Ireland; VTT Technical Research Center of Finland, Turku, Finland; Conway Institute of Biomolecular and Biomedical Research, University College Dublin, Ireland; Department of Health, Medicine and Care, Division of Diagnostics and Specialist Medicine, Linköping University, Linköping, Sweden; BioReperia AB, Linköping, Sweden; FINBB - Finnish Biobank Cooperative, Turku, Finland; Faculty of Medicine and Health Technology, Tampere University, Tampere, Finland; VTT Technical Research Centre of Finland Ltd., Espoo, Finland

**Keywords:** Neuroblastoma, MYCN, mitochondria

## Abstract

High-risk neuroblastoma is one of the most lethal childhood cancers. Half of these tumors are driven by MYCN gene amplification (MNA). Despite intensive chemo- and radiotherapy, only 40% of patients survive, and they often suffer from severe long-term side effects of these genotoxic treatments. Thus, new therapies are needed that are less toxic and more efficacious. Here, we identified diphenyleneiodonium (DPI) as a tool compound that preferentially targets MNA neuroblastoma. Using proteomic assays we investigated the DPI mode of action, finding that DPI induces the proteasomal degradation of MYCN and could reverse some alterations induced by high levels of MYCN. These include profound changes in the expression of proteins participating in the mitochondrial electron transport chain. Metabolic and biological assays suggested that alterations in mitochondrial function and the associated production of reactive oxygen species (ROS) are critical DPI targets in the context of MNA. DPI reduced the survival, and malignant transformation of neuroblastoma across a panel of cell lines at clinically achievable concentrations. DPI also shrank tumors and prevented metastatic spread in zebrafish models of MYCN-driven neuroblastoma. These findings suggest that processes impacted by complex I inhibitors could be valuable new targets for the development of non-genotoxic drugs against high-risk MNA neuroblastoma.

**Graphical abstract:** 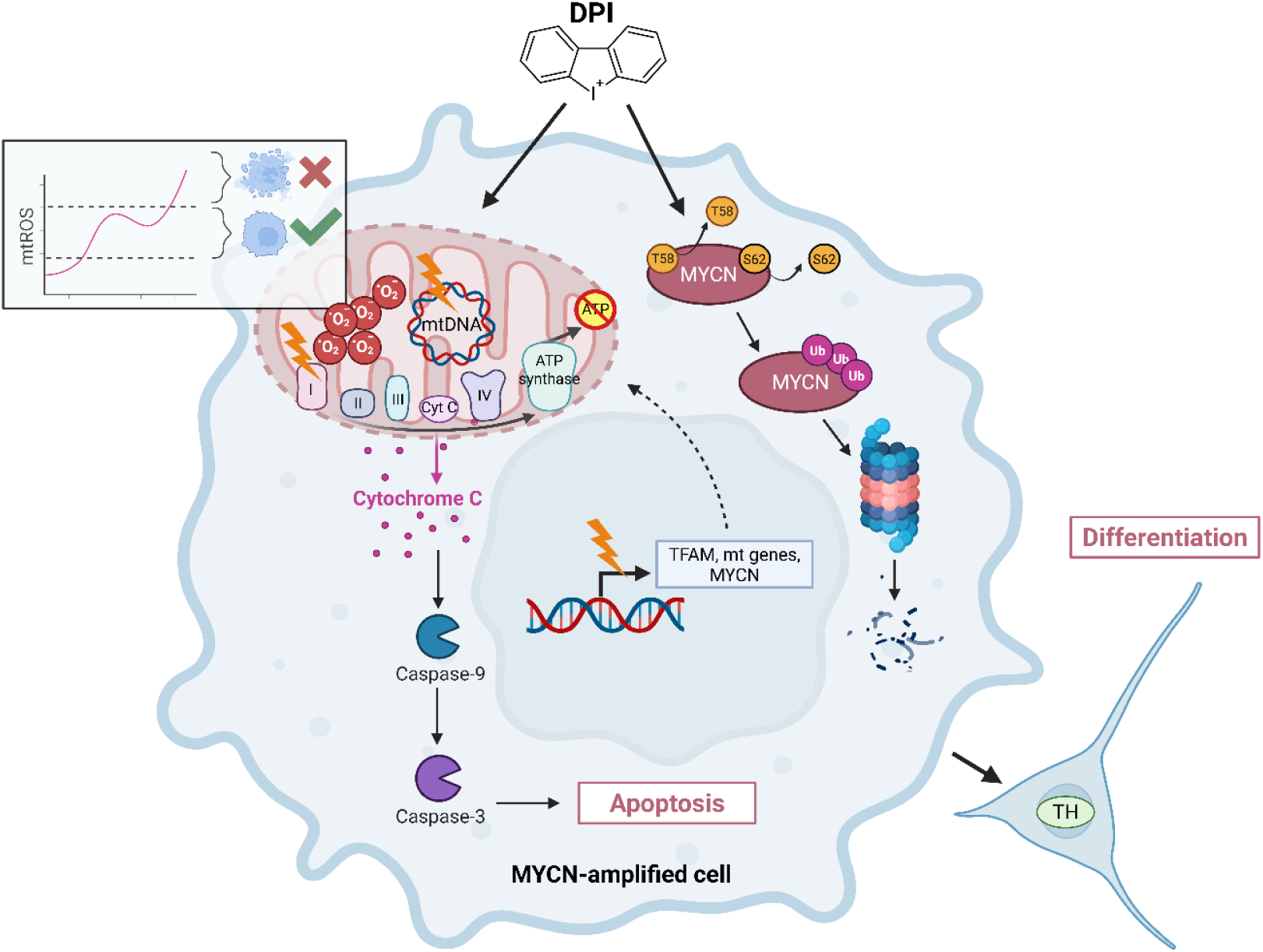

## Introduction

Neuroblastoma (NB) is a tumor of the peripheral sympathetic nervous system that accounts for only 6% of childhood cancer cases, but for 15% of deaths from childhood cancers[1]. This disproportionate risk is mainly due to high-risk NB that is driven by amplification of the MYCN gene. Patients with MYCN amplified (MNA) disease represent ca. 20% of all patients, and their 5-year event-free survival is only around 40%. They are usually treated with a combination of intensive genotoxic chemotherapy and radiation therapy. Nevertheless, many patients relapse, leaving them with limited or no treatment options. Furthermore, survivors may have severe long-term toxicities, including cardiovascular problems, hearing loss, secondary cancers, infertility, and endocrine deficiencies[2]. Thus, the current challenge is to identify therapies that are not genotoxic and more efficacious. This means that we must expand the drug target space and find compounds with new modes of action.

The obvious target is MYCN. It is a transcription factor with pleiotropic functions and considered as a master regulator of NB development[3]. However, MYCN is difficult to target directly as it lacks apparent pockets where drugs can bind to[4]. Thus, the current main strategies are to block critical MYCN functions and mechanisms that stabilize the inherently unstable MYN protein[5, 6]. However, so far none of these strategies have produced a clinically applicable approach.

Here, we used NB cell lines with regulatable MYCN expression to screen for compounds that preferentially inhibit cells with high MYCN levels. Amongst the most selective compounds targeting high MYCN expressing cells was diphenyleneiodonium chloride (DPI). This compound was originally described as NADPH oxidase inhibitor that reduces the generation of reactive oxygen species (ROS) in the mitochondria, but also has a broader range of targets including NAD/NADP-dependent enzymes in the tricarboxylic acid cycle and pentose phosphate pathway[7]. Newer studies showed that DPI also can induce ROS production by enhancing superoxide production from forward electron transport in complex I of the electron transport chain (ETC)[8]. Malignant cells, including NB cells, often produce more ROS than normal cells, as their increased demand for energy enhances oxidative phosphorylation (OXPHOS) and accompanying ROS production. This mechanism has been identified as a targetable vulnerability of MNA cells[9, 10]. Interestingly, MNA NB cells feature an enrichment of the ROS induced COSMIC mutational signature 18[11], suggesting that ROS contribute to the pathogenetic mechanism of NB.

Investigating the DPI mode of action in MNA cells, we found that DPI targets alterations in mitochondrial metabolism including OXPHOS and ROS production. Assessing DPI in both *in vitro* and in MYCN-driven zebrafish models of NB showed that DPI could efficiently counteract NB growth.

## Results

### DPI decreases viability more effectively in MNA cells and drives widespread proteomic alterations, particularly in mitochondrial functions

In order to screen for compounds that preferentially interfere with the viability of MYCN overexpressing NB, we used the SH-SY5Y cell line harboring a doxycycline inducible MYCN transgene (SY5Y/6TR (EU)/pTrex-Dest-30/MYCN cells; called SH-SY5Y/MYCN cells here)[12]. In these cells a 24-hour doxycycline treatment induces a 9-fold increase in MYCN protein expression (Fig. S1A). We screened 3,526 compounds (Fig. S1B) measuring cell viability. Using a cutoff of 3-standard deviations of the median, we identified 77 hits that selectively affected the viability of MYCN induced SH-SY5Y/MYCN cells (Table S1). These hits included compounds that were previously shown to have activity in NB, e.g., gitoxin, a cardenolide glycoside, which inhibits the growth of NB cell lines in mice[13]; inhibitors of MEK, JAK and PI3-kinases, which compromise cell survival[14]; estradiol, which promotes differentiation[12, 15]; BH3-mimetics, which induce apoptosis[16, 17]; histone deacetylase inhibitors, which are emerging as promising treatments for NB[18–20]; the inosine monophosphate dehydrogenase inhibitor mycophenolate mofetil, which inhibits proliferation and survival of NB cell lines[21]; the Aurora kinase inhibitor AT9283, which was tested in phase 1 clinical NB trials[22]; and inhibitors of DNA methylation (decitabine and 5-azacytidine), which sensitize NB cells to chemotherapeutics[23].

To select a candidate for further investigation, we focused on compounds that demonstrated effective cell viability inhibition at low nanomolar concentrations, since such potency could allow lower clinical dosing and improve the therapeutic window (Fig. S1C). We further prioritized compounds with minimal reported genotoxicity and low systemic toxicity. Based on these criteria, we selected diphenyleneiodonium chloride (DPI) for detailed follow-up studies (Fig. S1D). Moreover, DPI was previously reported to protect SH-SY5Y cells against ROS induced cell death[24, 25], suggesting that DPI may offer a selective treatment of MNA NB. Therefore, we examined DPI in more detail, especially in respect to its increased potency on MNA cells.

To assess the inhibitory effects of DPI on other NB cells, cell viability was measured in the MNA Be(2)-C, IMR-5/75, and IMR-32 cell lines. DPI potently reduced cell viability in all three MNA cell lines, with IC₅₀ values in the low nanomolar range (Fig. 1A). In addition, upregulating MYCN in SH-SY5Y/MYCN cells decreased the IC₅₀ approximately 1.6-fold, whereas downregulating MYCN in IMR-5/75 cells (+Dox) increased the DPI IC₅₀ more than 4-fold, indicating a MYCN-dependent contribution to DPI sensitivity, particularly pronounced in the MNA background (Fig. S1E-F).

**Figure 1.**
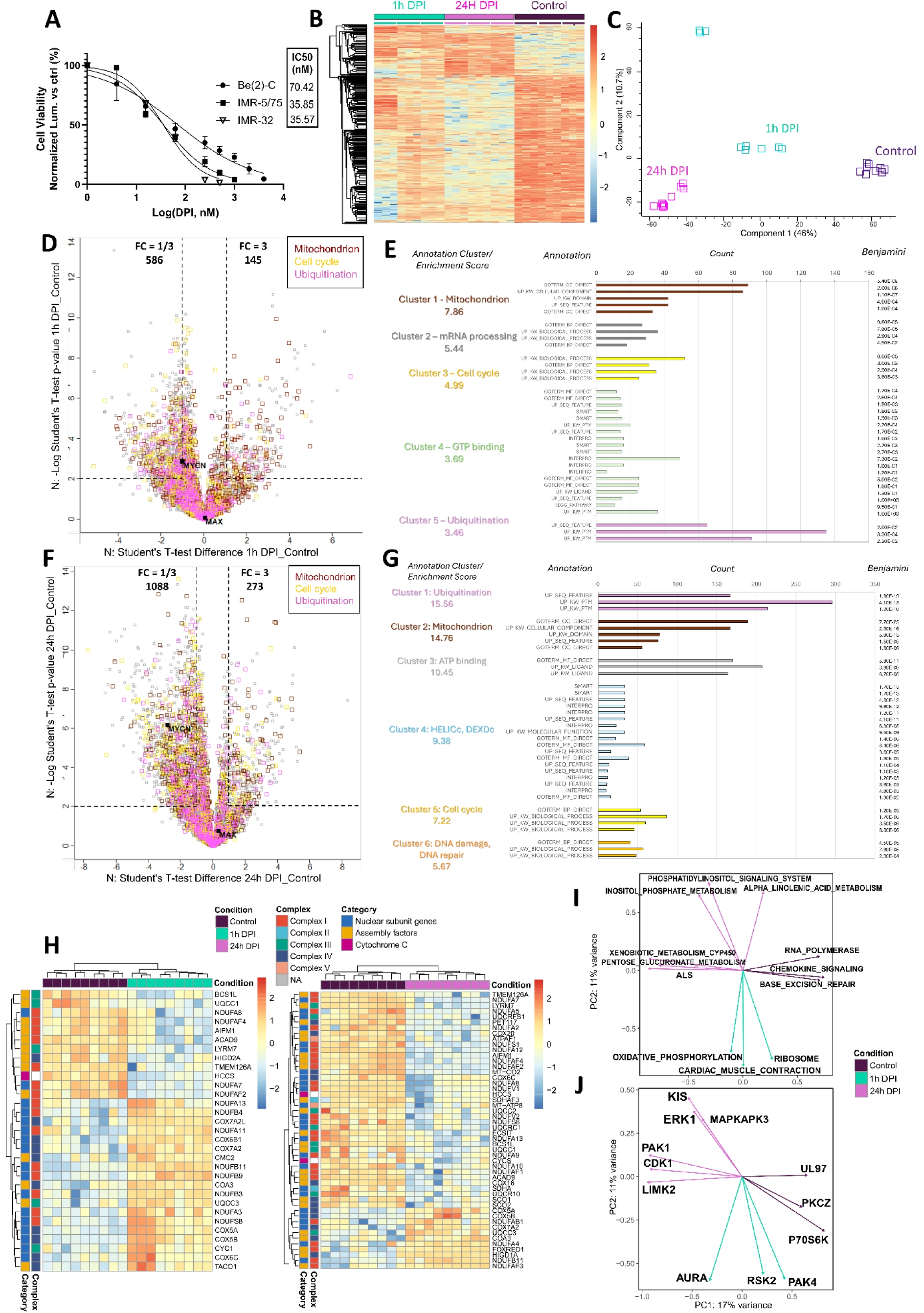
DPI decreases viability and triggers broad proteomic changes in MNA cells, notably affecting mitochondrial, cell cycle and ubiquitin-related protein networks. (A) Cell viability was measured as loss of ATP content using CellTiter-Glo in 3 MNA cell lines, after 72h treatment with serial dilutions of DPI (0 - 4000 nM). (B) Hierarchical clustering of 2,907 differentially expressed (T-test significant) proteins in control and DPI-treated (1h and 24h 10 μM DPI) Be(2)-C cells. Each condition has 3 biological and 3 technical replicates. The values in the heatmap represent the Z-transformation of log2(LFQ intensity). Red: high protein expression; blue: low protein expression (FDR < 0.01). (C) PCA analysis using Benjamini-Hochberg FDR = 0.01. (D) Scatter plot, 1h DPI versus untreated control. A 3-fold change (FC) was taken as cut-off. (E) The top 5 clusters out of 113 clusters following functional annotation clustering of up- and downregulated proteins using DAVID default annotation categories. (F) Scatter plot, 24h DPI versus untreated control. A 3-fold change (FC) was taken as cut-off. (G) The top 6 clusters out of 204 clusters following functional annotation clustering of up- and downregulated proteins using DAVID default annotation categories. (H) Hierarchical clustering of differentially expressed (T-test significant, FDR > 0.01) mitochondrial proteins in control and DPI-treated (1h and 24h) Be(2)-C cells grouped by respiratory complex affiliation and annotated as nuclear-encoded subunits or assembly factors proteins. Each condition consists of 9 columns: 3 biological replicates, each loaded 3x on the mass spectrometer. (I) Integration of the whole proteomic and phosphoproteomic data. Pathways from the protein expression profiling data contributing to the shared sources of variation in the data. (J) The kinases from the phosphoproteomic data contributing to the shared sources of variation in the data. Vectors are coloured according to the condition they are associated with. (Mean ± SEM, n = 3).

Given the higher MYCN levels of Be(2)-C cells compared to other MNA lines (Fig. S1G), we selected this model to probe DPI’s mechanism of action via mass spectrometry-based proteomic and phosphoproteomic profiling at 1h and 24h (Tables S2, S3). DPI profoundly altered protein expression (Table S2). Of the 5,042 quantifiable proteins 2,907 (∼57%) proteins were differentially expressed following DPI treatment (Fig. 1B). Principal component analysis (PCA) showed a clear separation of the three conditions (Fig. 1C). After 1 hour DPI treatment, 145 proteins were upregulated, and 586 proteins were downregulated threefold or more (Fig. 1D). Functional annotation clustering revealed that regulated proteins have functions related to mitochondria, mRNA processing, cell cycle, GDP-binding and ubiquitination (Fig. 1E). The GDP-binding cluster contains small G-proteins of the ARF, RAB and RAS families involved in vesicle transport and mitogenic signaling. After 24 hours DPI treatment, 273 and 1088 proteins were up- and downregulated, respectively (Fig. 1F). Cell cycle, mitochondria and ubiquitination remained the top altered functions. (Fig. 1G). Importantly, MYCN was downregulated by DPI treatment while the levels of MAX, a binding partner of MYCN remained unaffected.

Of the 3,860 quantifiable phosphosites, 1,384 were differentially abundant upon DPI treatment, following normalization to total proteome levels (Fig. S2A, Table S3). PCA analysis showed a clear separation of the three conditions (Fig. S2B). After 1h DPI treatment, 59 phosphorylation sites were upregulated and 28 were downregulated threefold or more (Fig. S2C). Functional annotation clustering mapped them to functions related to ubiquitination, RNA binding and mRNA processing (Fig. S2D). After 24 hours DPI treatment, 133 and 175 phosphorylation sites were up- and downregulated, respectively (Fig. S2E). Prominently altered functions included ubiquitination, nucleus, cell cycle, and translation initiation (Fig. S2F).

As the extensive changes in protein expression may influence the quantitation of phosphorylation sites, we integrated protein expression profiling with phosphoproteomics using multifactor analysis[26]. PCA analysis separated the conditions indicating that changes in both protein expression and phosphorylation independently contribute to the different molecular states induced by DPI treatment (Fig. S3A). The integrated data still discriminated the different treatment conditions revealing the shared sources of variation in the data represented by the factor loadings Z (Fig. S3B). The three conditions are mainly separated by Z1, while variations within conditions are mostly due to Z2. The biological processes and the phosphorylation changes associated with these variations were visualized by mapping the feature loadings onto pathways and kinases, respectively (Fig. 1I, J). These relationships provide information on the molecular mechanisms that mediate the DPI effects. For example, DPI treatment impacts oxidative phosphorylation and Aurora A (AURKA) kinase. Interestingly, in glioblastoma cells AURKA facilitates glycolysis, while suppressing OXPHOS[27]. DPI causes AURKA activation, albeit not significantly, (Fig. 1J, S3C), which could explain the similar effects of DPI on mitochondrial bioenergetics (Fig. 4B-D). The changes in inositol phosphate metabolism and phosphatidyl-signaling are correlated with increases in the activities of ERK1, KIS, and MAPKAPK3 (MK3) kinases after DPI treatment (Fig. 1I, J; S3C). ERK1 and KIS are involved in cell cycle progression, while MK3 promotes cell cycle arrest and senescence[28]. Pathways that characterize untreated control cells include pentose and glucuronate interconversion, which is involved in biosynthetic processes commonly upregulated in cancer[29], and amyotrophic lateral sclerosis, which is hallmarked by dysfunctional mitochondria[30]. Indeed, the corresponding kinases, PAK1, CDK1, and LIMK2 all have been associated with mitochondrial dysfunction[31–33]. Their activities were upregulated by DPI treatment (Fig. 1J, S3C). These results strongly suggested that mitochondria contain critical targets for the DPI effects.

### DPI induces MYCN degradation via phosphorylation-dependent proteasomal targeting

Ubiquitin-mediated protein degradation was among the most significantly enriched pathways in both total proteomic and phosphoproteomic analyses of MNA cells treated with DPI. Importantly, phosphorylation of MYCN at threonine 58 (Thr58) and serine 62 (Ser62) has previously been linked to its stability and degradation via the ubiquitin-proteasome system[34]. At 24 hours post-DPI treatment, phosphorylation at both sites was significantly decreased in the phosphoproteomic analysis (Fig. 2A), accompanied by a marked reduction in MYCN protein levels in the total proteomics data (Fig. 2B). These findings were validated by Western blotting (Fig. 2C). To determine whether reduced phosphorylation at these sites facilitates MYCN degradation via ubiquitination, we pre-treated cells with the proteasome inhibitor MG-132 prior to DPI exposure. MYCN protein levels were restored to baseline, indicating that the DPI-induced loss of MYCN is proteasome-dependent (Fig. 2D). We then performed a time-course analysis to assess whether MYCN loss was an early event following DPI treatment. Both MYCN protein and phosphorylation levels were reduced as early as 30 minutes after DPI addition, suggesting a rapid post-translational regulatory mechanism (Fig. 2E). These changes were evident by Western blot and phosphoproteomics (Fig. 2A). Finally, a dose-response experiment revealed that MYCN protein levels decreased in both MNA and non-MNA cell lines, with the MNA cells showing higher sensitivity at lower DPI concentrations (Fig. 2F).

**Figure 2.**
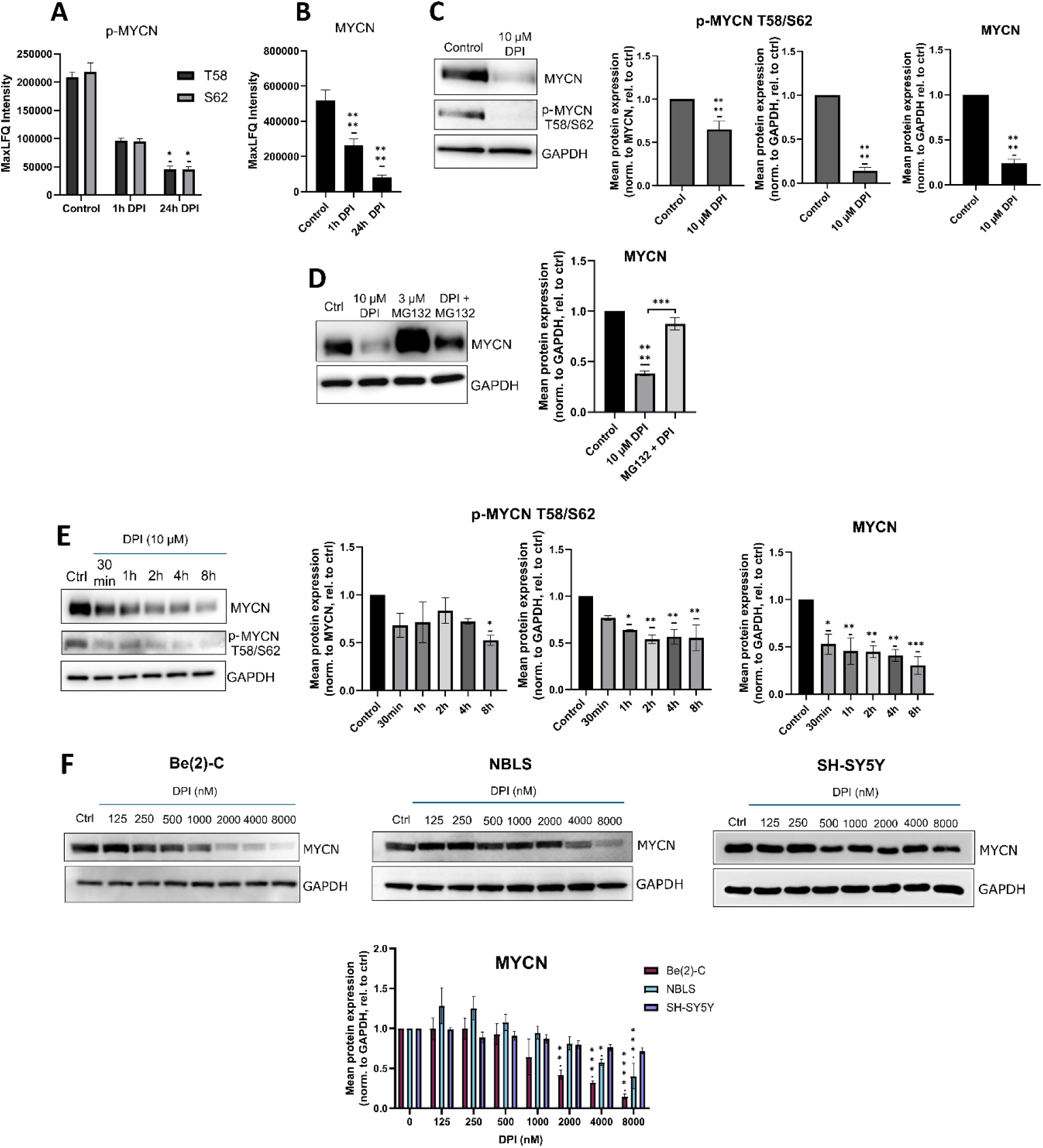
DPI affects MYCN phosphorylation status, impacting MYCN proteasomal degradation. **(A)** MYCN phosphorylation levels at Thr58 and Ser62 residues (p-MYCN) and **(B)** MYCN protein levels as measured by MS (Raw MaxLFQ Intensity) in Be(2)-C cells treated for 24h with 10 µM DPI (Significance indicated by stars is based on normalized data after statistical analysis). **(C–F)** Western blot analysis of p-MYCN (Thr58/Ser62) and/or total MYCN levels: **(C)** Validation of MS results in Be(2)-C cells treated with 10 µM DPI for 24 h. **(D)** MYCN levels following 4 h DPI treatment, after 1 h pre-treatment with 3 µM bortezomib. **(E)** Time-course treatment with 10 µM DPI for 30 min, 1 h, 2 h, 4 h, and 8 h. **(F)** Dose-response in Be(2)-C (MYCN-amplified), NBLS and SH-SY5Y (non-amplified) cells treated with serial DPI dilutions. GAPDH was used as a loading control for MYCN quantification. p-MYCN levels were normalized to both GAPDH and MYCN to confirm reproducibility. (Mean ± SEM, n = 3). Statistical significance was assessed by one-way ANOVA (p<0.05 = *, p<0.005 = **, p<0.0005 = ***, p<0.0001 = ****).

Together, these results suggest that DPI promotes early MYCN degradation through a mechanism involving reduced phosphorylation at regulatory residues and subsequent proteasomal targeting. This effect is observed even in cells with lower basal MYCN expression, indicating a shared vulnerability to DPI-induced destabilization of MYCN.

### DPI affects mitochondrial encoded gene expression and mitochondrial superoxide-mediated apoptosis of MNA NB cells

Our proteomics data in Be(2)-C cells showed that DPI alters the expression of ETC complex components (Fig. 1H). These changes involved proteins encoded by nuclear as well as mitochondrial DNA. To test whether DPI regulates gene expression on the transcriptional level to change the composition of ETC complexes, we quantified the expression of mitochondrial genes by RT-qPCR in a panel of DPI treated NB cells (Fig. 3A). DPI significantly reduced the expression of most genes in MNA cells, i.e., KCN, KCNR and Be(2)-C (Fig. S1G). This effect was attenuated or reversed in non-MNA NB cells (i.e., NBLS). Interestingly, DPI also downregulated the expression of MYCN mRNA in MNA cells (Fig. 3A).

**Figure 3.**
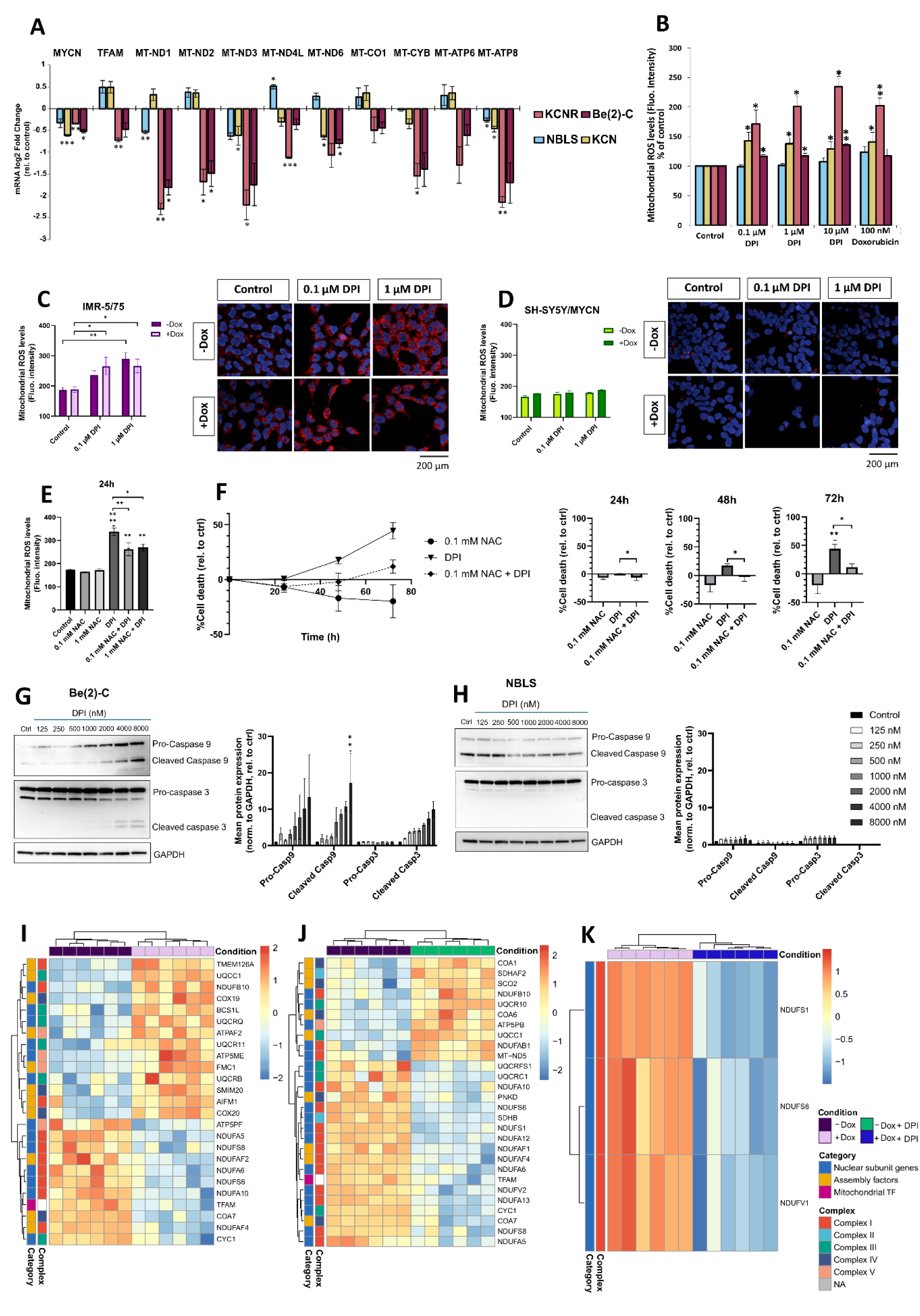
DPI targets mitochondrial gene expression and induces mtROS-dependent apoptosis in MYCN-amplified neuroblastoma cells. **(A)** RT-qPCR analysis of mitochondrial-encoded genes and MYCN in NBLS, KCN, KCNR, and Be(2)-C cells after DPI treatment (1 μM, 24h), shown as fold change over control. **(B)** Mitochondrial superoxide levels in NB cell lines treated with DPI for 24h, measured by MitoSOX Red-based flow cytometry. Doxorubicin (100 nM) used as positive control. **(C, D)** MitoSOX Red-based confocal imaging of mtROS in IMR-5/75 ±Dox **(C)** and SH-SY5Y/MYCN ±Dox **(D)** treated with DPI for 24h. **(E)** mtROS levels in Be(2)-C cells treated with NAC (0.1 or 1 mM) alone or 1 h prior to DPI (5 μM, 24h), measured by MitoSOX Red confocal imaging. **(F)** Cell death in Be(2)-C cells treated with NAC (0.1 or 1 mM) alone or 1h prior to DPI (5 μM, 24, 48, and 72h) measured with the CellTox Green indicator. **(G, H)** Western blot analysis of cleaved and pro-caspase-3 and -9 in Be(2)-C **(G)** and NBLS **(H)** cells after 24h DPI treatment (serial dilutions: 125–8000 nM). GAPDH was used as a loading control. **(I–K)** Hierarchical clustering of significantly altered mitochondrial proteins (T-test significant, FDR < 0.01) identified by MS-based proteomics in IMR-5/75: +Dox vs -Dox **(I)**, DPI-treated vs -Dox control **(J)**, and +Dox DPI-treated vs +Dox control **(K)**, organized by ETC complex and category. Each condition consists of 9 columns: 3 biological replicates, each loaded 3x on the mass spectrometer. (Mean ± SEM; n=3; *p < 0.05, **p < 0.01, ***p < 0.001, p<0.0001 = ****; Two-sample T-test, Ordinary One-Way ANOVA or Two-Way ANOVA).

DPI significantly reduced the expression of mitochondrial encoded genes, particularly those coding for complex I subunits (Fig. 3A). DPI was originally described as an NADPH oxidase inhibitor, but in addition to flavin containing enzymes may also inhibit NAD/NADP-dependent enzymes in the pentose phosphate pathway and tricarboxylic acid cycle[7]. DPI also can inhibit complex I in the ETC but may selectively enhance superoxide production from forward electron transport while blocking superoxide generation from reverse electron transport[8]. Superoxide dismutase decomposes the highly reactive superoxide to water and hydrogen peroxide which is further detoxified by antioxidant enzymes, such as peroxiredoxins, glutathione peroxidases, and catalases[35]. To selectively measure mitochondrial superoxide production upon DPI treatment we used the MitoSOX Red indicator (Fig. 3B). DPI selectively triggered superoxide production in the MNA cell lines, KCN, KCNR, and Be(2)-C, but not in the non-MNA NBLS cells. DPI induced superoxide to similar levels as doxorubicin, which is thought to exert cytotoxic effects in part via the generation of mtROS[36].

To further explore whether MYCN modulates mtROS production upon DPI treatment, we used two MYCN-regulatable NB cell models: the MNA IMR-5/75 line, in which endogenous MYCN expression can be downregulated by expression of a doxycycline inducible shRNA [37], and the non-amplified SH-SY5Y/MYCN line, where MYCN overexpression can be induced from a doxycycline inducible MYCN transgene. In IMR-5/75 cells, DPI significantly increased mitochondrial superoxide levels regardless of MYCN downregulation (Fig. 3C). In contrast, in SH-SY5Y/MYCN cells, DPI treatment did not augment mtROS levels, regardless whether MYCN expression was induced or not (Fig. 3D). Additionally, MYCN induction or repression alone did not significantly alter basal ROS levels in either model. These results suggest that high MYCN expression sensitizes cells to ROS induced cell death (see below).

When excessively accumulated, ROS can trigger apoptotic pathways[38]. To assess whether mtROS contribute to DPI-induced cell death, we pre-treated cells with the antioxidant N-acetyl-cysteine (NAC) before DPI exposure and monitored cell death over a 72-hour period (Fig. 3E). NAC treatment gradually rescued DPI-induced cell death in a time-dependent manner, with a significant decrease in cytotoxicity observed as early as 24 hours (Fig. 3F).

Inhibited mitochondrial respiration may lead to cytochrome C release from the mitochondrial intermembrane space to the cytosol, leading to activation of caspase-9 and caspase-3 and subsequent apoptosis[39]. DPI caused cytochrome C release in Be(2)-C but not in NBL-S cells (Fig. S4A). Cytochrome C release correlated with caspase-9 and caspase-3 cleavage and activation (Fig. 3G, H). Indeed, in Be(2)-C cells, DPI induced a significant accumulation of cleaved caspase-9, along with an increase in cleaved caspase-3 levels, which were not present in NBLS.

These findings suggest that MNA confers a mitochondrial vulnerability, consistent with prior evidence that c-MYC can regulate the expression of ETC components[40]. To investigate whether high MYCN levels impact the composition of ETC complexes, we performed total proteomics comparing IMR-5/75 cells under MNA (−Dox), downregulated MYCN (+Dox), and DPI-treated conditions (Figs. 3I–K, S4B-I, Table S4). MYCN downregulation (+Dox) affected the expression of 24 nuclear-encoded mitochondrial proteins (Fig. 3I). DPI treatment in −Dox cells altered 26 nuclear-encoded mitochondrial proteins and one mitochondrially encoded subunit (Fig. 3J). Notably, 10 of the proteins downregulated by DPI overlapped with those affected by MYCN downregulation, all regulated in the same direction. Moreover, both MYCN downregulation and DPI treatment downregulated TFAM and TFB1M, key mitochondrial transcription factors. In contrast, DPI treatment in +Dox cells led to significant changes in only three nuclear-encoded mitochondrial proteins, one of which was also affected by MYCN downregulation (Fig. 3K). Interestingly, ChIP-seq data from MNA NB cells (GSE183641) show that MYCN binds the promoter regions of NDUFAF4 and COA7 (Fig. S5). The expression of these proteins decreased ∼3-fold and ∼8-fold, respectively, upon MYCN knockdown, supporting their direct regulation by MYCN (Fig. 3I,J). The expression of these two proteins was also downregulated by DPI (Fig. 3I,J). Importantly, pathway enrichment analysis of DPI-treated IMR-5/75 -Dox cells revealed similar pathways to those observed in Be(2)-C cells (Figs. 1E, G; S4F, G), suggesting translational effects of DPI across different MNA cell lines.

Together, these data suggest that DPI disrupts mitochondrial-encoded gene expression in MNA cells, particularly complex I, creating a vulnerability to mtROS-induced apoptosis via cytochrome c release and caspase activation. This supports that DPI-induced cell death in MNA cells is at least partially mediated by ROS accumulation. While high MYCN expression appears to sensitize cells to ROS, acute MYCN up- or downregulation in the doxycycline regulatable cell lines does not significantly affect ROS generation (Fig. 3C,D) suggesting that ROS generation is not directly controlled by MYCN. However, an indirect effect is plausible. Our proteomic analysis in a MYCN-regulatable cell system indicates that high MYCN levels alter the composition of all ETC complexes (Fig. 3I). These changes may promote ROS production and may persist after acute regulation of MYCN expression. Interestingly, DPI can partially reverse the changes in ETC composition. This finding is also observable on a more global level, as pathways enriched upon MYCN downregulation closely mirror those affected by DPI treatment (Fig. S4C, E). Altogether, these findings support a MYCN-driven mitochondrial dependency that underlies the enhanced sensitivity of MNA cells to DPI.

### DPI modulates ATP production and promotes cell death, with MYCN amplification enhancing sensitivity

The substantial changes in the expression of ETC complex proteins mediated by MYCN and DPI likely affect mitochondrial functions. Therefore, we measured ATP production in the NB cells with regulatable MYCN using a Seahorse Analyzer (Fig. 4A-D). ATP can be produced by OXPHOS in the mitochondria or by glycolysis in the cytosol. Total ATP production was higher in IMR-5/75 than in SH-SY5Y/MYCN cells, but DPI reduced it in IMR-5/75 while slightly elevating it in SH-SY5Y/MYCN cells, although not in a significant manner (Fig. 4A). Interestingly, in IMR-5/75 more ATP (∼60%) was produced through OXPHOS than glycolysis in basal conditions, while the ratio was reversed in SH-SY5Y/MYCN cells (Fig. 4B-D). DPI robustly suppressed mitochondrial ATP production in IMR-5/75 and only non-significantly in SH-SY5Y/MYCN (Fig. 4C). Glycolytic ATP production was significantly enhanced by DPI in both cell lines, probably as a compensatory mechanism (Fig. 4D). Regulating MYCN levels only had a small, non-significant impact.

**Figure 4.**
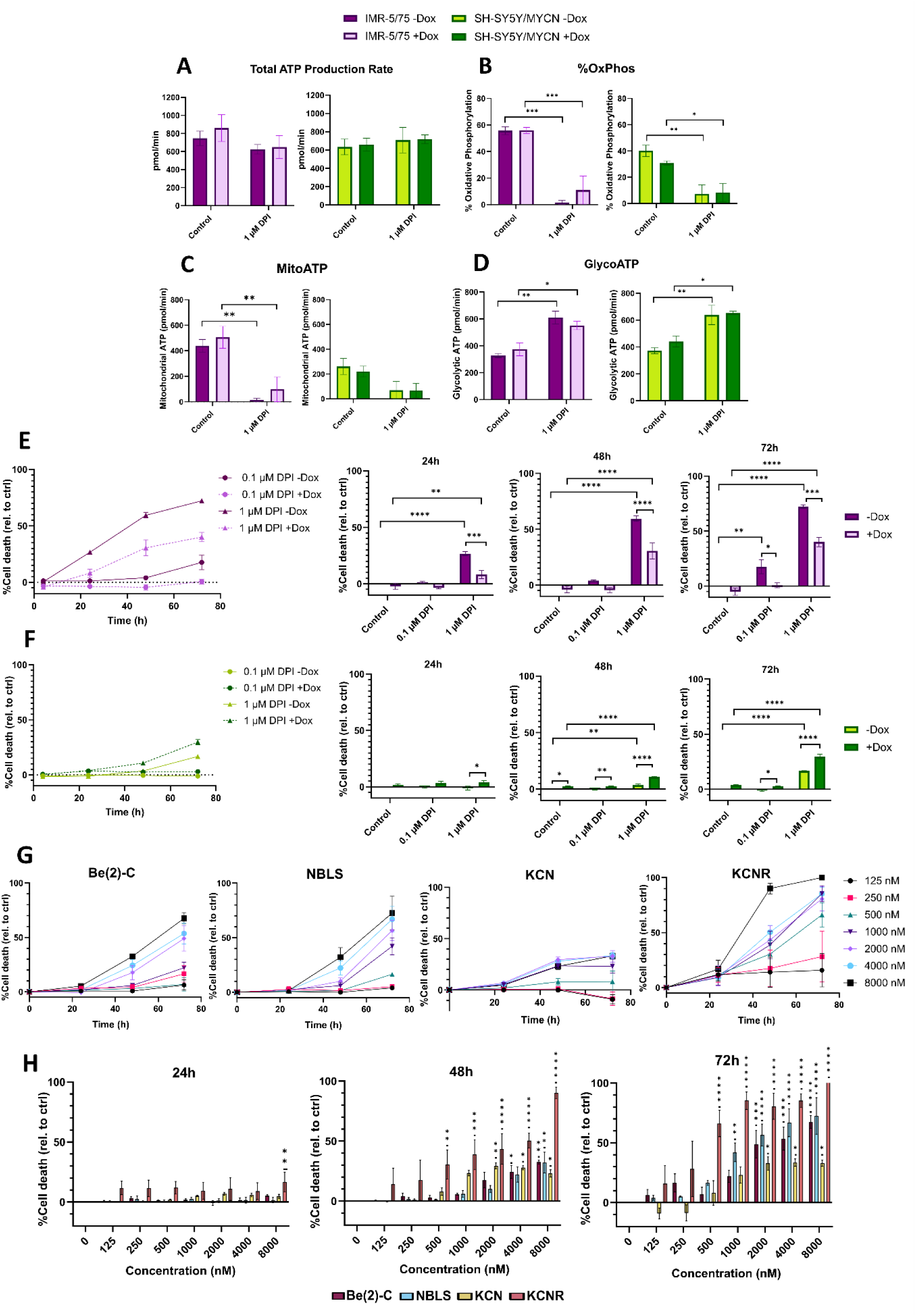
DPI reduces mitochondrial ATP production and promotes cell death in NB cells, with increased vulnerability in MNA cells. **(A)** Total ATP production rate, **(B)** percentage of ATP derived from OXPHOS, **(C)** mitochondrial ATP production, and **(D)** glycolytic ATP production measured in IMR-5/75 and SH-SY5Y/MYCN cells ±Dox and ±DPI (1 µM, 24h) using a Seahorse XFe24 Analyzer. **(E, F)** Cytotoxicity measured by CellTox Green (loss of membrane integrity) in IMR-5/75 **(E)** and SH-SY5Y/MYCN **(F)** cells treated with 0.1 or 1 µM DPI ±Dox over 72h. **(G, H)** Cell death assessed by CellTox Green in Be(2)-C, NBLS, KCN, and KCNR cells treated with DPI (0–8000 nM) over 72 h. (Mean ± SEM; n=3; *p < 0.05, **p < 0.01, ***p < 0.001, p<0.0001 = ****; Two-Way ANOVA).

To further characterize the impact of DPI in a MYCN-regulatable context, we assessed cytotoxicity. While total ATP production was not significantly affected by DPI after 24 hours (Fig. 4A), prolonged exposure led to marked reductions in cell viability, as measured by loss in ATP content, after 72 hours (Fig. S1E-F). A time-course analysis of cytotoxicity revealed effects as soon as 24 hours post-treatment, with increasing differences between MYCN-high and MYCN-low conditions at 48 and 72 hours in both cell lines (Fig. 4E, F). Notably, the MNA IMR-5/75 cells were overall more sensitive to DPI than the non-MNA SH-SY5Y/MYCN cells, and MYCN modulation significantly influenced the extent of cell death in both models. For instance, at 72 hours, 1 μM DPI induced ∼75% cell death in MYCN-high IMR-5/75 (-Dox) cells compared to ∼40% in MYCN-low (+Dox) cells (Fig. 4E, F).

To compare these effects across NB cell lines with distinct genetic backgrounds, we next assessed cell death kinetics in Be(2)-C, NBLS, KCN, and KCNR cells treated with serial dilutions of DPI (Fig. 4G, H). While non-MNA NBLS cells were also susceptible to DPI, they required higher concentrations and longer exposure times to induce comparable levels of cell death. Notably, the KCNR cell line, which is derived from a relapsed tumour after chemotherapy from the same patient as the KCN line, was most sensitive to DPI. This was particularly evident when compared to the treatment-naïve KCN cells, suggesting that treatment history and acquired vulnerabilities may enhance sensitivity to mitochondrial perturbation.

These results indicate that DPI selectively impairs mitochondrial ATP production in MNA IMR-5/75 cells, which rely more heavily on OXPHOS, while having limited impact on the more glycolytic SH-SY5Y/MYCN cells. The increased glycolytic ATP observed in both cell lines upon DPI treatment likely reflects a compensatory metabolic shift. This metabolic vulnerability is accompanied by enhanced sensitivity to DPI-induced cytotoxicity, particularly in MNA cells, supporting a MYCN-linked mitochondrial dependency that contributes to cell death.

### DPI counteracts the malignant phenotype of NB cells

The hallmarks of malignant transformation include resisting cell death and maintaining proliferation at the cost of differentiation[41]. MYCN promotes an undifferentiated, proliferative state in NB by actively repressing neuronal differentiation programs[42]. To assess the impact of DPI on these traits we carried out a soft agar colony forming assay and a differentiation experiment. Colony forming assays test the ability of single cells to survive and proliferate into a colony, which are typical properties of cancer cells[43]. The ability to form colonies decreased with lower MYCN expression levels, but DPI completely suppressed colony formation in all NB cells tested (Fig. S6A).

We further assessed differentiation through analysis of neuronal markers, neurite length and % of differentiating cells in IMR-5/75 and SH-SY5Y/MYCN cells ±Dox (Fig. 5A, B). For this, we compared DPI to retinoic acid (RA), which is a clinically used differentiation agent for neuroblastoma[44]. RA and even more DPI treatment significantly increased expression of the differentiation marker tyrosine hydroxylase (TH) after 5 days in IMR-5/75 cells, an effect reduced upon MYCN downregulation (+Dox) in DPI, but not RA treated cells (Fig. 5A). In SH-SY5Y/MYCN cells, DPI also moderately increased TH levels independently of MYCN induction. DPI treatment robustly decreased MYCN expression in IMR-5/75 cells and also in SH-SY5Y/MYCN when MYCN expression was induced. This observation is consistent with the notion that DPI has bigger effects when MYCN levels are higher. Morphological analysis revealed increased neurite length in IMR-5/75 cells treated with DPI, independent of MYCN levels (Fig. 5B). In contrast, RA increased neurite length significantly only when MYCN was knocked down. In SH-SY5Y/MYCN DPI has little effect, whereas RA only augmented neurite length when MYCN expression was induced, suggesting that RA has the best effects when MYCN is moderately overexpressed. This pattern was confirmed when quantifying the number of morphological differentiated cells, with the exception that DPI increased the % of differentiating cells solely in SH-SY5Y/MYCN cells upon MYCN overexpression, and that RA was more efficient than DPI.

**Figure 5.**
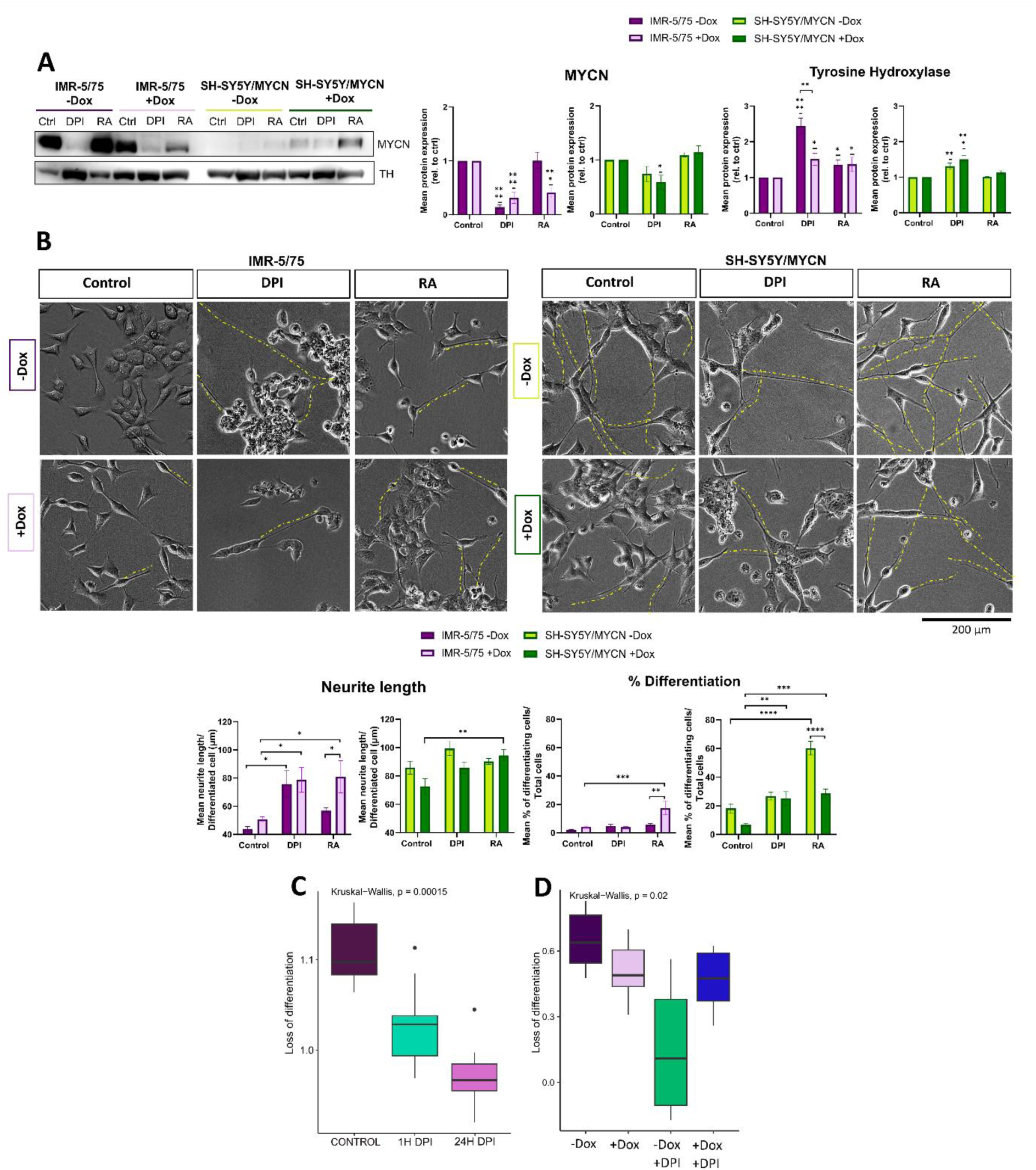
DPI induces differentiation of NB cells. IMR-5/75 and SH-SY5Y/MYCN cells were treated with -/+Dox and 0.5 µM DPI, 10 µM RA, or vehicle for 5 days. **(A)** Protein levels of the neuronal differentiation marker tyrosine hydroxylase (TH) and the pluripotency marker MYCN were assessed by Western blot. As these treatments disrupted traditional loading controls after 5 days, they could not be used for normalization. Equal loading was verified by BCA assay, and Coomassie Blue staining of membranes is shown in Supplementary Figure S6B as a control for total protein. **(B)** Neurite outgrowth and percentage of differentiating cells were quantified by microscopy. Cells with neurites ≥ 2 cell body diameters were considered differentiated. Three random fields per condition were analyzed from three independent experiments. Dotted yellow lines were used to outline neurite extensions indicative of neuronal differentiation. **(C, D)** Expression of a curated pluripotency-associated NB signature (BMI1, FN1, IGFBP2, MYCN, NES, NGFR, VIM) in **(C)** Be(2)-C cells treated with 10 µM DPI for 24 h and (D) IMR-5/75 -/+Dox cells treated with 0.5 µM DPI for 24 h based on MS-proteomic analysis. (Mean ± SEM; n=3; *p < 0.05, **p < 0.01, ***p < 0.001, p<0.0001 = ****; One-Way ANOVA on ranks; Two-Way ANOVA, Kruskal-Wallis test).

To complement these findings, we analysed proteomics data from DPI-treated Be(2)-C and IMR-5/75 -/+Dox cells using a curated NB pluripotency signature to assess whether DPI could shift cells from a pluripotent to a differentiated state[45, 46] (Figs. 5C, D). Indeed, DPI caused a loss of this signature, indicating a global shift towards differentiation. Interestingly, in IMR-5/75 cells, the analysis confirmed that DPI induced a more substantial loss of pluripotency in cells with higher MYCN levels (Fig. 5D). Together, these results suggest that DPI can counteract malignant properties of NB cells through both MYCN-dependent and independent mechanisms, with more pronounced effects when MYCN is increased (Fig. S6C).

### DPI inhibits NB growth and metastatic spread in zebrafish models of NB

To test whether DPI is also active against *in vivo* tumors, we used a well characterized zebrafish model of MYCN-driven NB. These fish overexpress an EGFP-MYCN transgene in the fish-equivalent of the adrenal gland and spontaneously develop NBs that faithfully recapitulate the genetic and molecular features of human MNA NB[47–51]. Administration of DPI (1 μM) to tumor bearing fish was non-toxic and resulted in a significant tumor regression after two weeks of treatment (Fig. 6A).

**Figure 6.**
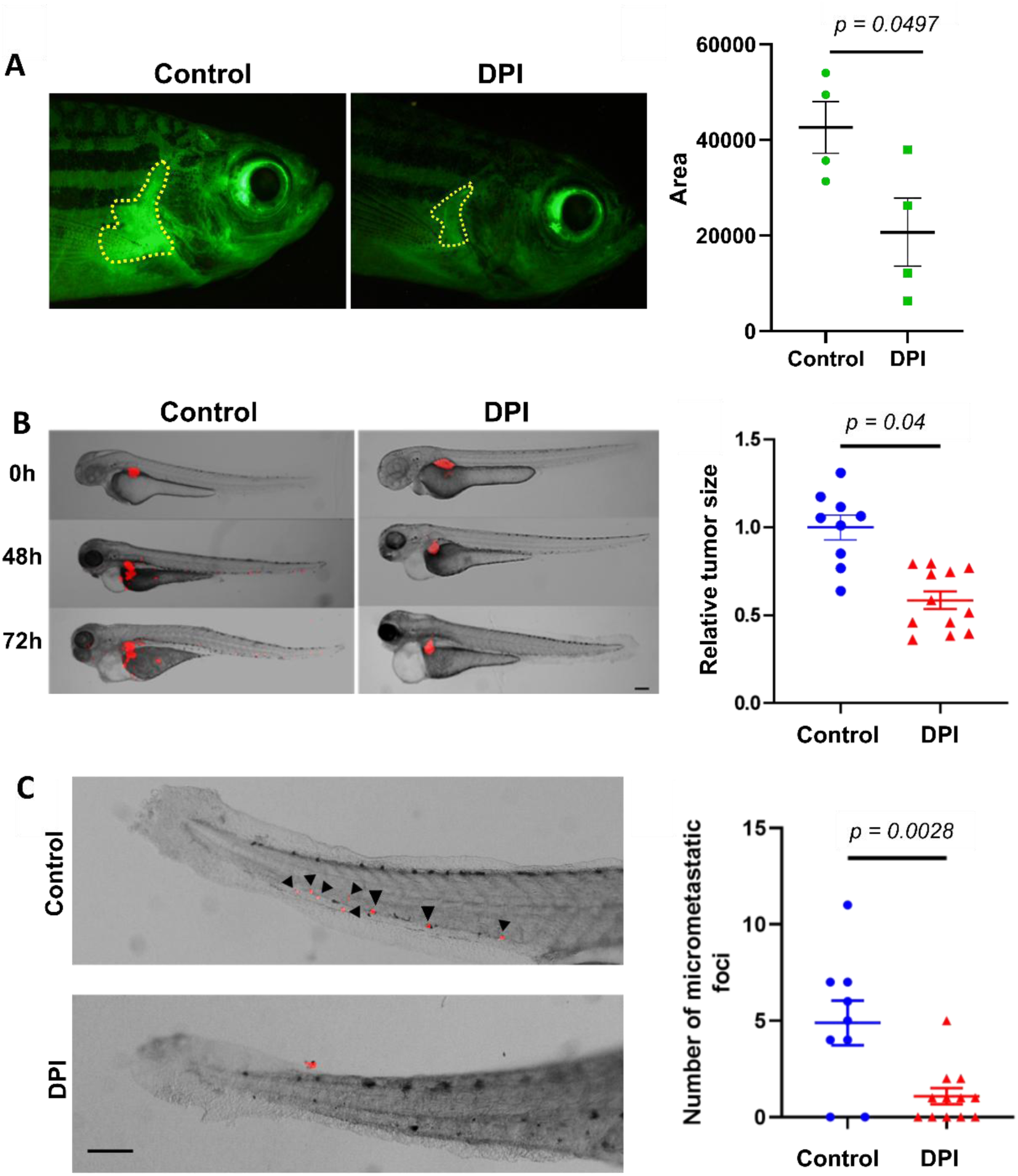
DPI suppresses tumor growth and metastatic spread in zebrafish models of MNA neuroblastoma. **(A)** Neuroblastoma growth in a zebrafish model driven by the overexpression of EGFP-MYCN. Tumor bearing fish (15 months of age) were treated with dimethyl sulfoxide (DMSO) control or DPI (1 µM, 14 days). Tumors (marked by broken yellow lines) are easily visualized due to EGFP-MYCN expression, and their sizes were determined by quantifying green fluorescence. (Mean ± SEM, Two-sample t-test, n = 4). **(B)** Be(2)-C cells labeled with red fluorescent dye were injected into the perivitelline space of 2 days post fertilization zebrafish embryos and treated with DPI (1 µM) for 72 hours. Tumor size was quantified by measuring red fluorescence. (Mean ± SEM, Two-sample t-test). **(C)** Black arrowheads indicate micro-metastatic foci in the tail of the fish. (Mean ± SEM, Two-sample t-test). Scalebars = 100 µm.

To assess whether tumors formed by human NB cells are also susceptible to DPI, we transplanted fluorescently labelled Be(2)-C cells into the perivitelline space of 2 days old zebrafish embryos and observed them over 72 hours (Fig. 6B, C). The transplants grew and spread throughout the fish. DPI treatment (1 µM) significantly reduced the primary tumor mass and number of micrometastases in the tail of zebrafish larvae compared to the non-treated group.

These findings show that DPI is effective against endogenously arising and transplanted NB suggesting that complex I inhibitors could be developed into a non-genotoxic treatment for NB.

## Discussion

Our results suggest that DPI is a promising tool compound for the development of new non-genotoxic NB treatments targeting complex I. Although we observed no toxicity in our zebrafish experiments, the use of DPI at micromolar concentrations may be limited by toxicity in mammalian species[52]. Therefore, to develop a clinically relevant drug, it is important to understand the mode of action, especially how DPI selectively targets biological processes specific to high-risk NB cells. While DPI was originally discovered as NADPH oxidase inhibitor, it transpired that it affects many more targets[7]. Our findings indicate that most of the relevant DPI actions are related to cells with MYCN gene amplification.

While MYCN is clearly a driver gene of high-risk NB, exploiting it as drug target has remained largely elusive[4]. MYCN itself seems not druggable as it has no obvious hydrophobic pockets where small molecule inhibitors can bind. Therefore, indirect approaches have prevailed. For example, Aurora A (AURA) kinase binds to MYCN and protects it from degradation. Aurora A kinase inhibitors (AURAi) destabilize this interaction causing MYCN degradation[53]. They showed promising results in preclinical models but proved ineffective in clinical trials in humans[54]. Recent improvements combined AURAi with ATR kinase inhibitors, exploiting the fact that MYCN activates AURA, and AURA inhibition induces DNA damage that requires ATR kinase for repair[55]. Thus, inhibiting both AURA and ATR prevents MYCN from limiting DNA damage. This mechanism is interesting, as according to our proteomics data DNA repair is an enriched pathway, and DPI treatment reduced the expression of ATR to 29% within 24 hours. Aurora A levels also diminished, albeit to a lesser extent, to 64%. These changes could plausibly contribute to both the decrease in MYCN protein and the enhanced apoptosis in response to DPI.

DPI also modulates the phosphorylation sites of MYCN that regulate its activation and subsequent degradation, namely T58 and S62[56]. Mitogenic kinases (CDK1, ERK1/2, mTOR) phosphorylate S62 enhancing MYCN activity, while the subsequent phosphorylation of T58 induces S62 dephosphorylation and MYCN degradation. Our normalized phosphoproteomics data show that DPI decreases the phosphorylation of MYCN at T58 and S62. In addition, DPI also reduces the phosphorylation of S371 and S375 after 24 hours. The corresponding phosphorylation sites in c-MYC, S344 and S348, inhibit both the transcriptional and transforming activity of c-MYC[57]. Thus, DPI may control MYCN activity and stability by multiple mechanisms. Interestingly, the extent of MYCN reduction by DPI correlated with MYCN status suggesting that the suppressive mechanism is under control of MYCN itself. Our previous results showed that MYCN regulates the expression of its own interaction partners[12], and analyzing the impact of DPI on the expression of MYCN interaction partners could provide further insights into how DPI preferentially impacts overexpressed MYCN. Interestingly, we found that MAX expression remained unchanged upon DPI treatment, suggesting that MYCN and MAX might be regulated independently under specific stress conditions. Using MYCN-regulatable cell lines and a panel of NB cell lines with diverse MYCN status and genetic backgrounds, we showed that DPI preferentially targets cells with MNA but also affects non-MNA cells at higher doses and longer exposure times. This highlights DPI’s broader therapeutic potential.

Apart from these effects on MYCN itself, DPI overlaps with MYCN in affecting the function of mitochondria. MNA fundamentally changes the metabolic functions of mitochondria[58] and renders MNA cells dependent on MYCN to maintain the function of these altered mitochondria[58]. Our proteomics profiling revealed profound changes in the expression of mitochondrial ETC proteins, which likely would modify the composition and function of ETC complexes. Amplified MYCN can increase both glycolysis and OXPHOS[58]. Using Seahorse analysis, we found that MNA cells preferentially use mitochondrial ATP generated through OXPHOS. DPI profoundly suppresses OXPHOS while triggering a compensatory increase in glycolysis, possibly leading to metabolic imbalance. This metabolic shift likely represents an adaptive response, yet it is insufficient to prevent DPI-induced cytotoxicity.

As DPI can inhibit ETC complex I in the OXPHOS chain leading to the production of superoxide[8], the generation of ROS induced by DPI in MNA cells is likely closely related to this metabolic imbalance. DPI was reported to scavenge ROS[24, 25], but the altered composition of ETC complexes in MNA cells may subvert this protective function into a deleterious liability. In our experiments, DPI induced superoxide production, caspase-dependent apoptosis, cytotoxicity, differentiation, and decreased proliferation and MYCN levels more strongly in MNA cells. While DPI also reduced cell viability and increased cell death in non-MNA cells, this required higher concentrations and longer exposure times. Although ROS levels were not directly modulated by MYCN regulation alone, compensatory mechanisms may buffer this effect. However, under DPI-induced complex I stress, these compensatory mechanisms might become overwhelmed, pushing ROS beyond a critical threshold that triggers cell death particularly in MNA cells. In our experiments the ability of DPI to induce superoxide production, apoptosis, cytotoxicity and decrease of proliferation was linked to higher MYCN levels. The only biological DPI effect that was MYCN independent, was the induction of differentiation. Thus, it seems that the most relevant targets of DPI that suppress MNA NB reside in the mitochondria exerting the metabolic imbalance and superoxide production induced by MYCN amplification.

Although mitochondria are moving into the limelight as targets for new cancer drugs[59], a wealth of previous experience shows that monotherapies are unlikely to succeed in the clinic. Therefore, complex I inhibitors will need to be combined with other drugs. Our results suggest that DPI synergizes with chemotherapeutic drugs currently used for NB treatment, such as vincristine, and also with agents that modulate mitochondrial ROS production[60]. Taken together, these data indicate that the MYCN dependent metabolic alterations are the most relevant DPI target for further drug development.

## Methods

### Cell lines

Cells were generous gifts from J Schulte (Charite, Berlin, Germany) and F Westermann (DKFZ, Heidelberg, Germany). SH-SY5Y and NBLS (single MYCN gene copy) and KCN, KCNR, Be(2)-C (MNA) cells were cultured in RPMI-1640 (Gibco), supplemented with 10% fetal bovine serum (Gibco), 2 mmol/L L-glutamine (Gibco), and 1% penicillin-streptomycin solution (Gibco). For SY5Y/6TR (EU)/pTrex-Dest-30/MYCN cells (here called SH-SY5Y/MYCN cells), where MYCN overexpression is doxycycline inducible[12], the above culture media was supplemented with Blasticidin (7.5 µg/mL) and G418 (200 µg/mL). For the MYCN-amplified IMR-5/75 cells, where MYCN is downregulatable by doxycycline induction of an shRNA against MYCN[37], the culture media also contained Blasticidin (5 µg/mL) and Zeocin (50 µg/mL). Doxycycline was used at 1 µg/mL concentration.

### Drug screen

SH-SY5Y/MYCN cells were seeded in 384-well plates (1500 cells per well) and treated with 1μg/ml Doxycycline (Sigma) for 24 hours, which resulted in a 11-fold increase in MYCN mRNA. Then, uninduced and Doxycycline induced cells were treated with the following drugs: Biomol (80 known kinase and phosphatase inhibitors; at 10, 1, 0.1 and 0.01 µmol/L), LOPAC (1280 compounds; at 1 and 0.1 µmol/L), Microsource Cancer (80 compounds) and Microsource Spectrum (2000 compounds including most of the known drugs and other bioactive compounds and natural products; 1 and 0.1 µmol/L) totaling 3500 compounds (Table S1). Viability was measured 76 hours later using the CellTiter-GLo Luminescence Cell Viability Assay as described previously[12].

### Chemicals

Diphenyleneiodonium chloride (DPI) was purchased from Enzo Life Sciences (#BML-CN240-0010). Dimethyl sulfoxide (DMSO, vehicle) (#D2650), MG-132 (#M7449) and retinoic acid (#R2625) were from Merck. Doxorubicin was purchased from Selleck Chemicals (#S1225).

### Western blotting

Cells were lysed with lysis buffer (1% Triton X, 150 mM NaCl, 1 mM MgCl_2_, and 20 mM Tris–HCl, pH 7.5) supplemented with Complete Mini (Roche) protease and PhosStop (Roche) phosphatase inhibitors. Proteins were resolved in 10% NuPAGE Bis-Tris Mini gels (Life technologies) at 100 V and transferred to a polyvinylidene difluoride membrane (Invitrolon) at 30 V, 70 min. Membranes were blocked using 5% non-fat dried milk (Sigma) in TBS with 0.05% Tween (TBS-T) for 1 h and incubated overnight at 4°C with primary antibodies against MYCN (#sc-53993, Santa Cruz), p-MYCN Thr58/Ser62 (#9401, Cell Signaling), GADPH (#2118, Cell Signaling), cytochrome C (#Ab110325, Abcam), caspase 3 (#9665, Cell Signaling), caspase 9 (#9508, Cell Signaling), tyrosine hydroxylase (#2792, Cell Signaling) diluted in 1:1000 in 5% BSA/TBS-T. Membranes were then washed with TBS-T and incubated for 1 h at room temperature with peroxidase conjugated anti-mouse or anti-rabbit IgG (1:5000 dilution, 7076 and 7074, Cell Signaling). Blots were developed using Advanced Molecular Image Imager and ChemoStar Software. Quantification of blots was achieved using ImageJ software v1.44p[61].

### Cell fate assays

Cell viability and cytotoxicity measurements were performed using CellTiterGlo and CellTox Green assays (both from Promega) according to the manufacturer’s instructions. For differentiation, cells were observed using light microscopy and images were taken at 10x, 20x, 40x magnification using an Olympus 2-620 camera connected to Olympus CKX41 microscope. Neurite elongation was measured using the NeuronJ plugin from ImageJ software[61]. Cells with neurite length/cell body width ratios ≥2 were considered differentiated. Neurite length was calculated by dividing mean neurite length/differentiated cell; and % differentiation was calculated by dividing the mean % of differentiating cells/total cells in each field of view.

For the soft agar assay, 1 ml of 0.8% base agarose layer was placed in each well of 6-well plates, then plates were cooled for 5 min at 4°C to solidify agarose. Then 1 ml of upper agarose layer (0.48%) with 10000 cells was added on the solidified base layer. Then plates were cooled for 5 min, then 1 ml of growth media (+/- 1 µM DPI) was added to each well. Plates were incubated at 37°C and 5% CO_2_ for 20 days, monitoring for colony formation. Media (+/- DPI) was replaced every 4 days. Colonies were stained by adding PBS containing 4% formaldehyde and 0.005% crystal violet to each well for 1 hour.

### Proteomics and phosphoproteomics

Cells were seeded in 15cm^2^ dishes in 20 ml culture media. Upon reaching 70-80% confluence, Be(2)-C were treated with 10 µM DPI for 0, 1 and 24 hours in reverse order, in triplicates. IMR-5/75 cells were seeded under the same conditions. MYCN expression was downregulated by treating the cells with 1 µg/mL doxycycline. Following MYCN downregulation, cells were treated with 0.5 µM DPI for 24 hours, in triplicates. Cells were lysed in ice cold 8M urea/50 mM Tris-HCL pH 8.0, supplemented with phosphatase and protease inhibitors (Roche). Samples were then sonicated for 2 X 9 seconds at 10% power using Branson Sonifier® 250 connected to Syclon Ultrasonic Homogenizer. To reduce protein samples, 8 mM dithithreitol (dTT) was added to the samples and incubated in a thermomixer for 60 minutes at 30°C. Then, samples were alkylated by adding 20 mM iodoacetamide and incubated in a thermomixer for 30 minutes at 30°C in the dark. Samples were diluted in 50mM ammonia bicarbonate. 10 µL of sequencing grade trypsin were added into samples and left overnight on a thermomixer at 37°C to allow sample digestion. The following day, formic acid was added to 1% final concentration to stop sample digestion.

The Glygen TopTipTM C18 protocol (200µL) was utilized for sample clean up. Samples were centrifuged at 5000 rpm for 5 minutes to remove particulates. 200 µL samples were added into a 1.5mL tube and centrifuged at 3000 rpm for 2 minutes. 200 µL equilibration and washing buffer (0.1% Trifluoracetic acid (TFA) in high performance liquid chromatography (HPLC) grade water were added and the mixtures were centrifuged at 2500 rpm for 2 minutes. Supernatants were dispensed to waste. 200 µL wetting and elution buffer (80% Acetronitrile (ACN) in 0.1% TFA) were added to the pellets and centrifuged at 3000 rpm for 2 minutes. Samples were evaporated for approximately 30 minutes at 30°C in an Eppendorf Concentrator.

#### TiO_2_ (titanium dioxide) phosphopeptide enrichment

Dried peptide samples were re-suspended in 500 µL Binding Buffer. TiO_2_ ratio of beads to sample was weighed out at 10:1. 1 mg of TiO_2_ was re-suspended in 5 µL of Binding Buffer and mixed on a shaker for 5 minutes at room temperature. 5 µL of TiO_2_/Binding Buffer was added to each sample. Supernatants were removed and kept in -80°C. TiO_2_ beads were washed with 1 mL Wash Buffer A (80% Acetonitrile, 1% TFA) and centrifuged at 5000 rpm for 1 minute. Wash Buffer A was then removed, and this step was repeated thrice. Next, TiO_2_ beads were washed with 1mL Wash Buffer B (80% Acetonitrile, 0.1% TFA) then centrifuged at 5000 rpm for 1 minute. Wash Buffer B was then removed. 50 µL of Elution Buffer (50% Acetonitrile, 25% NH4OH) were added 2 times to elute the phosphopeptides. Samples were dried for approximately 20 minutes in an Eppendorf Concentrator at 30°C.

#### LC-MS/MS Method (Bruker timsTof Pro)

After trypsin digestion and sample clean up using C18 TT2 TopTips (Glygen), samples were run on a Bruker timsTof Pro mass spectrometer connected to a Bruker nanoElute nano-lc chromatography system. Tryptic peptides were resuspended in 0.1% formic acid. Each sample was loaded onto an Aurora UHPLC column (25cm x 75μm ID, C18, 1.6 μm) (Ionopticks) and separated with an increasing acetonitrile gradient over 90 minutes at a flow rate of 250 nl/min at 45°C.

The mass spectrometer was operated in positive ion mode with a capillary voltage of 1500 V, dry gas flow of 3 l/min and a dry temperature of 180 °C. All data was acquired with the instrument operating in trapped ion mobility spectrometry (TIMS) mode. Trapped ions were selected for ms/ms using parallel accumulation serial fragmentation (PASEF). A scan range of (100-1700 m/z) was performed at a rate of 10 PASEF MS/MS frames to 1 MS scan with a cycle time of 1.16s.

#### Data analysis

The raw data was searched against the Homo sapiens subset of the Uniprot Swissprot database (reviewed) using the search engine Fragpipe using specific parameters for trapped ion mobility spectra data dependent acquisition (TIMS DDA). Each peptide used for protein identification met specific Fragpipe parameters, i.e., only peptide scores that corresponded to a false discovery rate (FDR) of 0.01 were accepted from the Fragpipe database search. The normalised protein intensity of each identified protein was used for label free quantitation (LFQ).

Data pre-processing and statistical analysis was performed using Perseus (version 2.1.2.0)[63] for Figs. 1, 3 and S2 and S3. Phosphoproteomics results were normalized to total proteomics to account for protein abundance. Significant proteins were displayed in a heatmap (FDR < 0.01 for whole proteomics and FDR < 0.05 for phosphoproteomics). Selected proteins were further analysed using DAVID for pathway enrichment[64].

### Bioinformatic analysis

The Fragpipe proteinGroups.txt files for both the whole- and phospho-proteomics were filtered to remove known contaminants and protein groups containing no protein. LFQ intensity values were log base 2 transformed, with zero values being assigned NaN to circumvent undefined values. Filtering was applied to keep only those proteins/phosphopeptides with at least 6/9 valid values in at least one drug treatment condition. NaN values were then reassigned to zero before median centering. Site- and condition-specific imputation was performed using the scImpute function in PhosR[65] if the protein was quantified in more than 50% of samples from the same drug treatment. Sample-wise tail-based imputation was implemented with the tImpute function[65]. Linear regression models were used to determine the relationship between a protein’s expression and its phosphorylation levels. The residuals from the linear models were used to normalize the phosphoproteomics data by the whole proteomics data. Differentially abundant phosphosites and proteins between control and DPI-treated samples were identified using the Limma R package. An FDR threshold of 0.05 was used to determine statistical significance.

### Multiset correlation and factor analysis

Kinase-substrate relationships were obtained from the PhosphoSitePlus database (https://www.phosphosite.org/homeAction.action), while pathway-gene relationships from the KEGG database were accessed through MSigDB. Both the pathways and kinases were scored in the whole proteomics and phosphoproteomics datasets using the GSVA function in R. The resulting scores, which represent the activity levels of kinases and pathways in each sample, were then used as input for Multiset Correlation and Factor Analysis[26]. This analysis was conducted using default parameters to identify shared factors of variation across both omics’ datasets. Kinases and pathways that contributed most to the variation between sample conditions were selected based on their absolute feature loadings.

### ChIP-seq data analysis of MYCN binding sites

ChIP-seq data from the MYCN-amplified neuroblastoma cell line IMR-32 (GEO accession GSE183641) were analyzed using the R2 ChIP-seq Genome Browser (http://r2.amc.nl), focusing on MYCN occupancy at promoter regions. Signal intensity profiles were compared between siControl (IMR32_27_siCtrl MYCN), input (IMR32_25_siCtrl Input) and siMYCN (IMR32_15_siMYCN)-treated conditions to identify differential binding.

### Cell fractionation

Mitochondria were isolated using the Qproteome Mitochondria Isolation Kit according to manufacturer’s instructions (Qiagen). Ten million cells have been used for the isolation of mitochondria and the purity of the fraction was assayed by Western blotting for cytosol (GADPH) or mitochondria (cytochrome C) specific proteins.

### MitoSOX Red staining

Mitochondrial superoxide levels were assessed using the MitoSOX Red fluorescent indicator (Thermo Scientific) by flow cytometry and live cell imaging, according to manufacturer’s instructions.

Flow cytometry: 1x10^5^ cells were seeded in a 12 well plate and treated with vehicle (DMSO) or DPI for 24h. After treatment, 1 ml of HBSS buffer with 2.5 μM of MitoSOX™ reagent was added to the cells and incubated for 10 minutes at 37°C. Cells were detached using Versene solution and after two washes in HBBS were re-suspended in 500 ml of HBBS for flow cytometry analysis. Forward and side scatter gates were established to exclude non-viable cells and cell debris from the analysis. Auto-fluorescence signals generated by unlabeled cells were used as negative controls in each experiment. Flow cytometric analysis was performed on an Accuri C6 instrument by using the FL2 detector (Optical Filter 585/40 nm) and analyzed with CFlow® Software (Accuri, Ann Arbor, MI, USA). The data were expressed as median fluorescence intensity of three independent experiments.

For imaging, cells were typically seeded in 100 μL of RPMI phenol red-free media per well in duplicate on a black 96-well plate with a clear bottom (Thermo Scientific). To ensure IMR-5/75 and SH-SY5Y/MYCN cells remained attached, the plates were pre-coated with 100 μg/mL Poly-D-Lysine (Sigma-Aldrich). 24 hours post-doxycycline induction, the cells were treated with 100 μL of DPI or vehicle and incubated at 37°C. Following the designated incubation period, cells were rinsed once with DPBS containing Ca^2+^/Mg^2+^ (Invitrogen). MitoSOX Red dye (4 μM) was then added along with the Hoechst 33342 nuclear counterstain (1 μg/mL) (Thermo Scientific) in DPBS containing Ca 2+ /Mg 2+ and incubated for 20 minutes at 37°C, shielded from light. After incubation, cells were washed twice, and fresh RPMI phenol red-free media was added before live-cell imaging. The MitoSOX Red indicator was detected using the Texas Red filter (Ex/Em: 510/580 nm), while Hoechst was detected using the HOECHST 33342 filter (Ex/Em: 353/483 nm).

### Live cell imaging

Confocal images were captured across four optical planes spaced 1 μm apart (ranging from -1.0 μm to 2.0 μm) within 25 x 25 fields of view, with a 10% overlap per well, using an Opera Phenix High Content Screening System (PerkinElmer). The imaging was performed with either a 40x or 63x/1.15 NA water-immersion objective under live cell conditions (37 °C and 5% CO2). The images obtained were analyzed using an optimized pipeline tailored for each cell line on Harmony® v.4.9 high-content analysis software (PerkinElmer). The software’s building blocks were employed to identify and calculate the mean dye parameters per well, including the number of nuclei and MitoSOX Red mean intensity per cell.

### RT-qPCR

Total RNA was isolated from NB cell lysates using the RNeasy kit (Qiagen, UK). Reverse transcription was carried out on 1000 ng of total RNA using Superscipt III reverse transcriptase (Invitrogene) according to the manufacturers’ instructions. Quantification of gene expression by real-time PCR was performed on an ABI Prism 7700 Sequence Detection System, (Applied Biosystems Inc., UK). MT-ND1, MT-ND2, MT-ND3, MT-ND4L, MT-ND6, MT-ATP6, MT-ATP8, MT-CO1, MT-CYB, TFAM, and MYCN gene expression was examined using specific Taqman gene expression assays (Applied Biosystems Inc., UK). GADPH was used as endogenous control in the assay.

### Metabolic measurements

Metabolic activity was evaluated using a Seahorse XF24 Analyzer (Seahorse Biosciences) through an ATP rate test. IMR-5/75 and SH-SY5Y/MYCN cells were seeded at 37,500 cells per well in PDL-coated XF microplates and incubated in XF RPMI medium with glucose, sodium pyruvate, and glutamine. After six hours, doxycycline was added to induce gene expression in specific wells. On the second day, cells were treated with DPI or vehicle, and the XFe24 Sensor Cartridge was hydrated overnight. On the third day, cells were washed with assay medium, incubated, and the ATP rate test was conducted by sequentially injecting oligomycin A and a mix of rotenone and antimycin A. Post-assay, cells were fixed with glutaraldehyde, stained with Crystal Violet for normalization, and absorbance was measured. Data was analyzed using Wave software to generate key metabolic parameters, including Oxygen Consumption Rate (OCR), Proton Efflux Rate (PER), and Extracellular Acidification Rate (ECAR), followed by statistical analysis in GraphPad Prism.

***Zebrafish experiments*** were approved by the University College Dublin Animal Research Ethics Committee and the Healthcare Products Regulatory Authority (AE18982/P038). zebrafish (Danio rerio) were maintained at 28°C on a 14-hour light/10-hour dark cycle. Xenografts: At 2dpf, dechorionated zebrafish embryos were microinjected with fluorescently labelled Be(2)-C cells. Prior to microinjection, embryos were transferred to prewarmed 2% agarose gel and anaesthetized with 0.05mg/mL Tricaine. Be(2)-C cells were fluorescently labelled with DiI. Approximately 100-500 cells were injected into the perivitelline space of each embryo. Following injection, embryos were screened and randomly assigned a treatment arm. Each embryo was placed in an individual well of 24 well plate containing 600uL PTU ± DPI (1 uM). Embryos were incubated at 35 °C for duration of treatment (72 h). Embryos were imaged daily using fluorescent microscopy (Nikon Eclipse C1) to monitor tumour progression and metastasis. Data analysis was carried out using ImageJ, Excel and GraphPad Prism.

The Tg[dbh:MYCN-EGFP] transgenic zebrafish was a generous gift from A. T. Look (Dana-Farber Cancer Institute, Harvard Medical School)[51]. Adult Tg[dbh:MYCN-EGFP] zebrafish were monitored for fluorescent tumor masses biweekly. Zebrafish developing the fish-equivalent of human neuroblastoma in the interrenal gland (age, 15 months; n = 4) were treated every 48 hours with DPI for 2 weeks. Variance in tumor size for all fish at pretreatment was <5%. Zebrafish were anesthetized by 0.016% tricaine before microscopic analysis. An Olympus SZX10 fluorescence stereomicroscope equipped with an Olympus DP71 camera was used to capture images. The fluorescent region posterior to the gills, which corresponds to the region of neuroblastoma growth[51], was analyzed using ImageJ software (version 1.44p).

## Acknowledgements

This work received funding from Science Foundation Ireland and National Children’s Research Centre / Children’s Health Ireland through the Precision Oncology Ireland grant 18/SPP/3522, the Research Ireland Frontiers for the Future Program grant 22/FFP-A/10729, the Irish Research Council grant GOIPG/2020/1361, the Pathological Society of Great Britain and Ireland, Horizon 2020 programme grant H2020-RISE/GA734907 (3D-NEONET), European Union FP7 grant 259348-2 “ASSET,” and financial support of Children’s Health Foundation under the management of Science Foundation Ireland under the Frontiers for the Future Programme Grant Number 21/FFP-P/10130. Mass spectrometry was performed at The Comprehensive Molecular Analytical Platform which was funded by The Research Ireland Research Infrastructure Programme (18/RI/5702). This project was partially supported by the Fight Kids Cancer Funding Programme, supported by Imagine for Margo, KickCancer, Fondatioun Kriibskrank Kanner, Federazione Italiana Associazioni Genitori e Guariti Oncoematologia Pediatrica (FIAGOP), and Cris Cancer Foundation.

## Author contributions

Conceptualization, W.K. and M.H.; Investigation, S.E., St.M., A.A., Si.M., K.I., K.W., M.H.; Formal Analysis, S.E., St.M., K.I., M.H., D.E.; Writing - all authors contributed to writing the manuscript; Visualization, S.E., D.E., M.H.; Funding Acquisition, W.K., M.H., L.J., St.M., A.A.; Resources, K.I., K.W., L.J.; Supervision, L.J., M.H., and W.K.

