## Supplementary Figures for "Diphenyleneiodonium chloride inhibits MYCN-amplified neuroblastoma by targeting MYCN induced mitochondrial alterations"

**A**

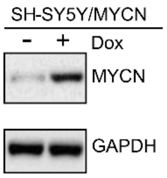


**B**

| Compound library | Compounds | Concentrations (µM) |
|---|---|---|
| Sigma Lopac | 1280 | 0.1; 1 |
| Microsource Spectrum | 2000 | 0.1; 1 |
| Microsource Cancer | 80 | 0.01; 0.1; 1; 10 |
| BioMol, kinase & phosphatase inhibitors | 84 | 0.01; 0.1; 1; 10 |
| Individually ordered compounds | 82 | 0.1; 1; 10 |
| Total compounds | 3526 | |

**C**

| Compound library | Name | Final concentration | SY5Y-NMYC | SY5Y | Reported side effects | Target |
|---|---|---|---|---|---|---|
| BMI | 5-IODOTUBERCIDIN | 250 nM | -0.395419374 | -0.18211 | Limited genotoxicity, potential cytotoxicity | Adenosine metabolism |
| Selleck | ABT-263(Navitoclax) | 926 nM | -0.165790957 | -0.00101 | Thrombocytopenia, Lymphopenia, Testicular toxicity | BCL-2, Mitochondrial apoptotic pathway |
| MS | AKLAVINE HYDROCHLORIDE | 250 nM | -0.315745667 | -0.19928 | Unclear genotoxicity, some cytotoxicity | Antimicrobial, Antioneoplastic |
| MS | ALBENDAZOLE | 250 nM | -0.415215021 | -0.21027 | Low genotoxicity, cytotoxicity at high doses | Anthelmintic; binds tubulin, disrupting microtubules in parasites |
| MS | ALPINETIN METHYL ETHER | 250 nM | -0.658795651 | 0.020035 | Natural compound, little data | Anti-inflammatory, anti -cancer properties |
| Selleck | Axitinib | 926 nM | -0.153124634 | -0.06295 | No clear genotoxicity, some cytotoxicities reported | Tyrosine kinase inhibitor (VEGFR 1/2/3 inhibitor; anti-angiogenesis) |
| MS | beta-CARYOPHYLLENE ALCOHOL | 250 nM | -0.531499392 | -0.05389 | Natural compound, low cytotoxicity | Sesquiterpene with anti -inflammatory and analgesic properties |
| Selleck | BIBW2992 (Tovok) | 926 nM | -0.152897207 | -0.04718 | Low genotoxicity, some cytotoxicity | Irreversible EGFR and HER2 tyrosine kinase inhibitor |
| MS | BICUCULLINE (+) GABA | 250 nM | -0.619143259 | 0.125612 | Neurotoxicity at high doses | GABA_A receptor antagonist |
| Selleck | Decitabine | 103 nM | -0.195364659 | -0.03542 | Genotoxic and cytotoxic | DNA methyltransferase inhibitor |
| LOPAC | Diphenyleneiodonium chloride | 250 nM | -0.327620412 | -0.13822 | Some reported off -targets effects but safe at low doses | NADPH oxidase and flavoprotein inhibitor, Mitochondrial inhibitor |
| LOPAC | Emetine dihydrochloride hydrate | 250 nM | -0.433602364 | 0.072963 | Cytotoxic, potential cardiotoxicity | Protein synthesis inhibitor |
| Selleck | Etoposide (Etopophos) | 926 nM | -0.200272818 | -0.12377 | Genotoxic and cytotoxic | Topoisomerase II inhibitor |
| Selleck | Fenbendazole | 926 nM | -0.180667705 | -0.04913 | Low toxicity | Anthelmintic; binds tubulin, disrupts microtubules in parasites |
| LOPAC | Flunarizine dihydrochloride | 250 nM | -0.327161801 | 0.059049 | Generally well tolerated, some mild side effects reported | Calcium channel blocker |
| MS | LATHOSTEROL | 250 nM | -1.413639673 | -0.04085 | Cholesterol precursor | Cholesterol biosynthesis intermediate |
| Selleck | LY2228820 | 103 nM | -0.171965191 | -0.12269 | Limited genotoxicity/cytotoxicity data | p38 MAP kinase inhibitor/EGFR inhibitor |
| Selleck | Methotrexate | 308 nM | -0.15575862 | -0.00412 | Genotoxic and cytotoxic | Dihydrofolate reductase inhibito |
| Selleck | Mubritinib | 103 nM | -0.178439161 | -0.0744 | Limited genotoxicity/cytotoxicity data | ErbB2/HER2 inhibitor, Mitochondrial inhibitor |
| MS | PACLITAXEL | 250 nM | -3.517317371 | -0.35879 | Genotoxic and cytotoxic | Stabilizes microtubules, prevents mitosis |
| BMI | STAUROSPORINE | 25 nM | -0.279459835 | -0.1245 | Highly cytotoxic, no clinical use due to toxicity | Broad kinase inhibitor, Mitochondrial apoptotic pathway |
| MS | THIRAM | 250 nM | -0.369485778 | -0.34826 | Some toxicity | Antifungal and pesticide, inhibits cellular enzymes |
| Selleck | Tretinoin | 103 nM | -0.174119431 | -0.14213 | No genotoxicity, midly cytotoxic, used clinically for the treatment of NB | Nuclear retinoic acid receptors; induces differentiation |

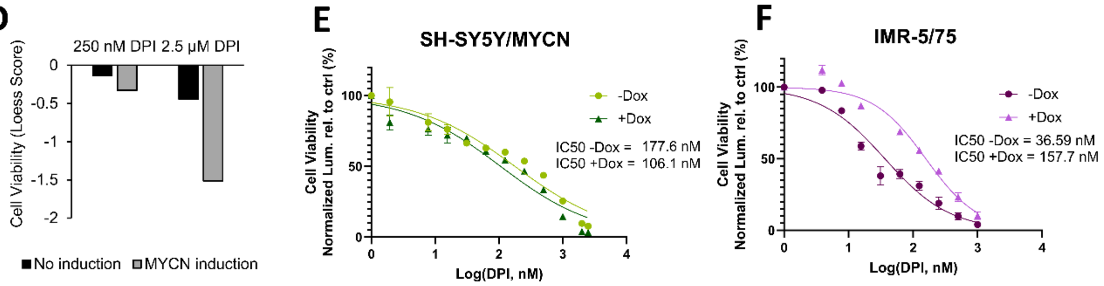


**G**

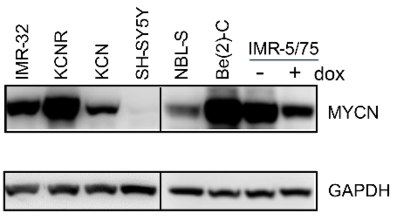


***Supplementary Figure S1. Drug screen identifies DPI as a potential MYCN-selective hit.*** **(A)** MYCN protein levels in SH-SY5Y/MYCN cells before and after a 24 hour induction with doxycycline (dox). MYCN protein expression was normalized to GAPDH. **(B)** Compound libraries used for screening. **(C)** Compounds shown were selected from the 77 hits based on their ability to inhibit cell viability in SH-SY5Y/MYCN Dox+ cells ('MYCN high') at low nanomolar (nM) ranges. **(D)** Cell viability assessed by CellTiter-Glo (ATP content) 250 nM or 2500 nM DPI treatment after 72 h in SH-SY5Y/MYCN ±Dox (drug screen, n=1). **(E,F)** Cell viability assessed by CellTiter-Glo after 72 h DPI treatment (0 – 2000 nM) in **(E)** SH-SY5Y/MYCN ±Dox, and **(F)** IMR-5/75 ±Dox. **(G)** Western blot of MYCN expression in the indicated cell lines. MYCN protein expression was normalized to GAPDH.

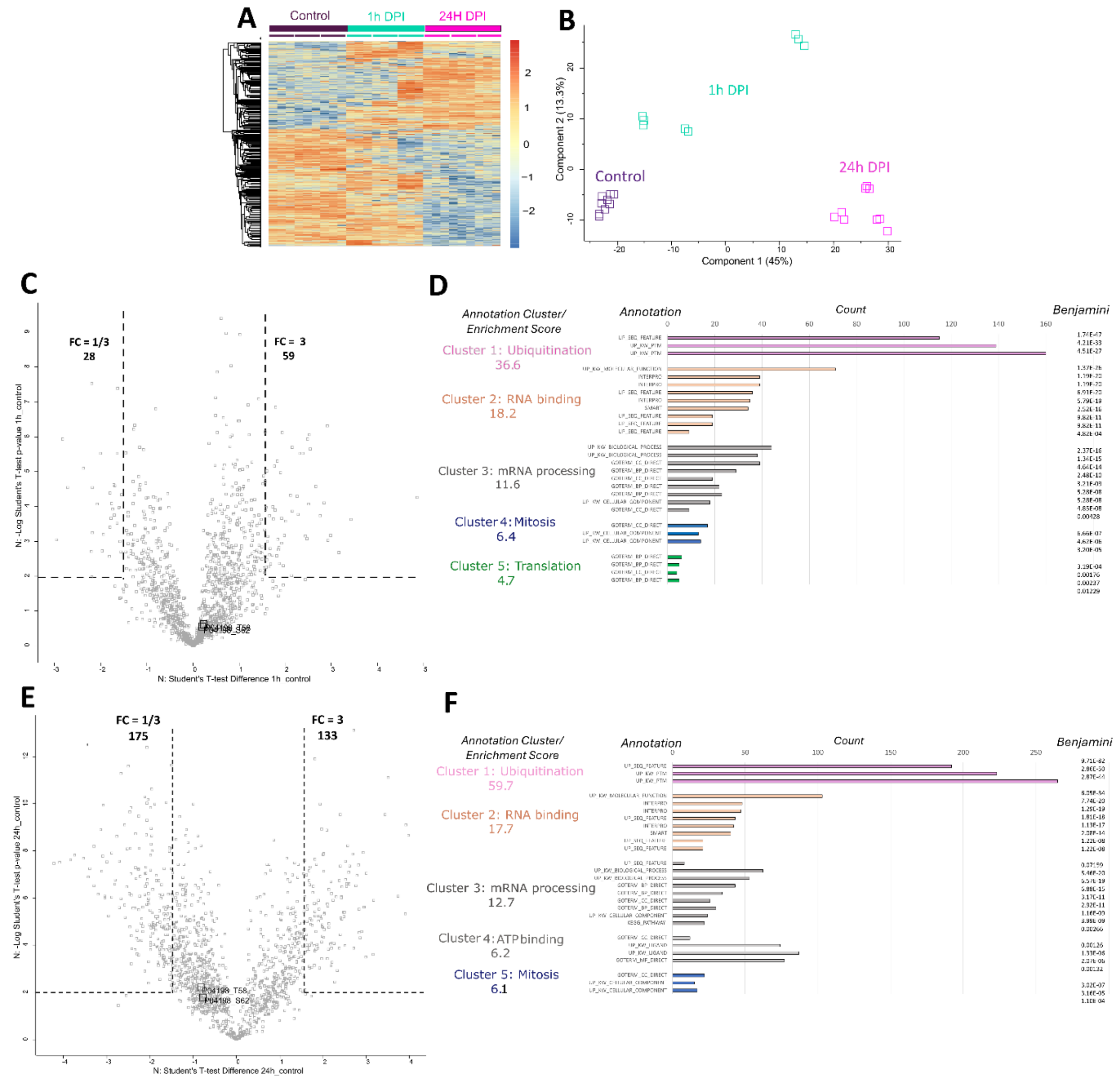


***Supplementary Figure S2. Phosphoproteomics of Be(2)-C cells treated with DPI.*** **(A)** Hierarchical clustering of 1,384 differentially abundant (t-test significant) phosphorylation sites in control and DPI-treated (1h and 24h 10 µM DPI) Be(2)-C cells. Each condition has 3 biological and 3 technical replicates. The values in the heatmap represent the Z-transformation of log2(LFQ intensity). Red represents high protein expression while blue represents low protein expression. Permutation-based FDR < 0.05. **(B)** PCA analysis using a Benjamini-Hochberg FDR = 0.05. **(C)** Scatter plot, 1h DPI versus untreated control. A 3-fold change (FC) was taken as cut-off. **(D)** The top 3 clusters out of 42 clusters following functional annotation clustering of up- and downregulated proteins using DAVID default annotation categories. **(E)** Scatter plot, 24h DPI versus untreated control. A 3-fold change (FC) was taken as cut-off. **(F)** The top 5 clusters out of 204 clusters following functional annotation clustering of up- and downregulated proteins using DAVID default annotation categories.

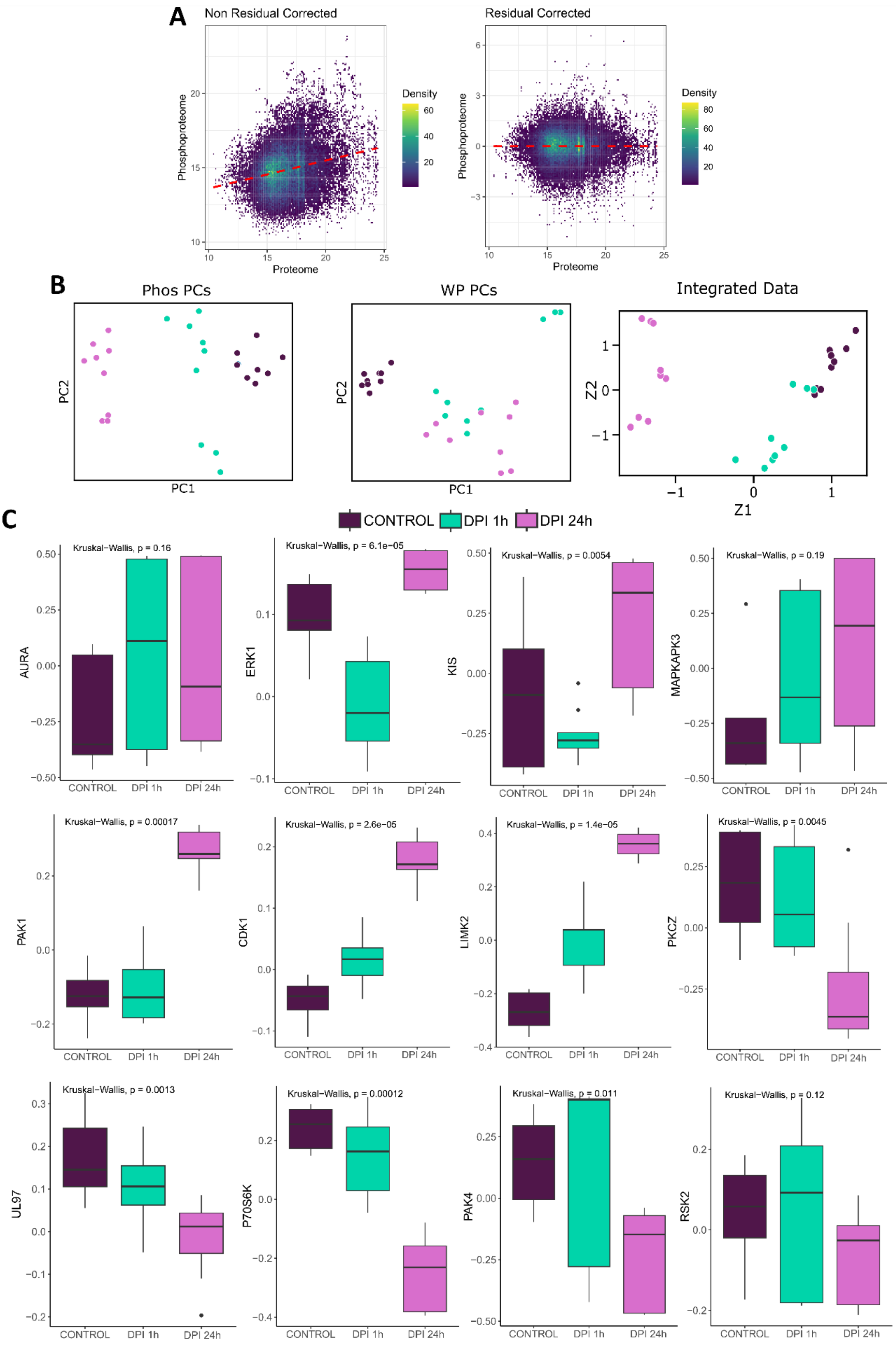


***Supplementary Figure S3. Integration of total proteomics and phosphoproteomics data of DPI-treated Be(2)-C cells by multiset correlation and factor analysis.*** **(A)** The relationship between the expression of a protein and its phosphorylation sites. Left panel: prenormalization of the phosphoproteomics by the whole proteomics data. Right panel: post-normalization of the phosphoproteomics by the whole proteomics data. **(B)** PCA analyses of data frames that represent the activity of pathways and kinases in each sample (Left and middle image). Integration of the whole proteomic and phosphoproteomic data (right). Z1 and Z2 represent shared sources of variation in the data. **(C)** Kinase activity changes inferred by Kinase–Substrate Enrichment Analysis (KSEA). The kinases represented account for the largest proportion of variability observed in the dataset.

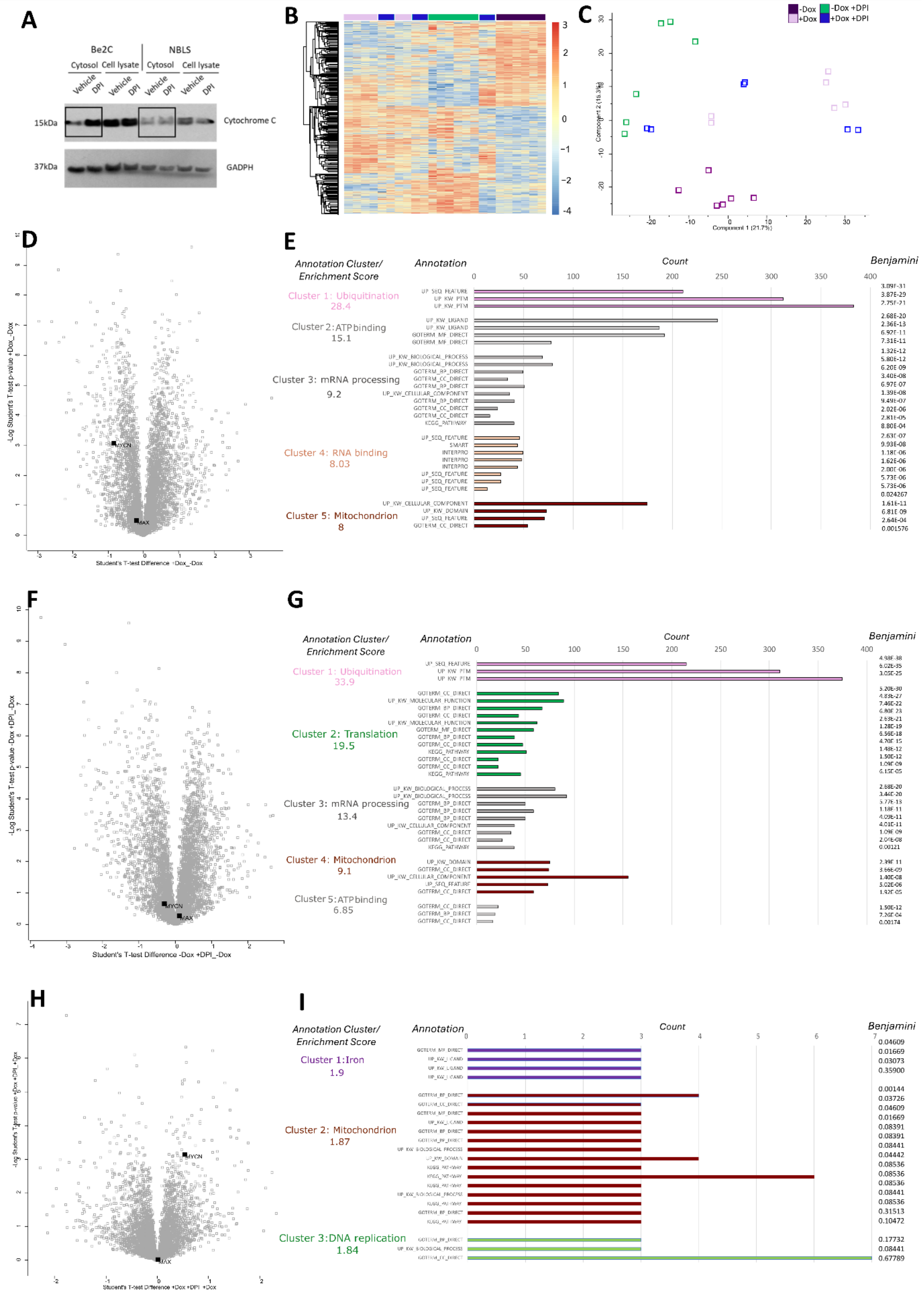

***Supplementary Figure S4. Cytochrome c release and mitochondrial proteomic remodeling following DPI treatment.*** **(A)** Cytochrome c release was analysed by measuring cytochrome c levels in the cytosolic fraction and in the total lysates of NBLS and Be(2)-C cells upon 24 h

DPI treatment (1 µM) by Western blotting. (Mean ± SEM, n = 3). **(B)** Hierarchical clustering of 2,278 differentially expressed (T-test significant) proteins in -Dox, +Dox, -Dox +DPI and +Dox +DPI treated IMR-5/75 cells (0.5 µM DPI treatment for 24 h). Each condition has 3 biological and 2 technical replicates. The values in the heatmap represent the Z-transformation of log2 (LFQ intensity). Red: high protein expression; blue: low protein expression (FDR < 0.01). **(C)** PCA analysis using Benjamini-Hochberg FDR = 0.01. **(D)** Scatter plot, +Dox versus -Dox untreated control. **(E)** The top 5 clusters out of 245 clusters following functional annotation clustering of up- and downregulated proteins using DAVID default annotation categories. **(F)** Scatter plot, 24h DPI versus -Dox untreated control. **(G)** The top 5 clusters out of 234 clusters following functional annotation clustering of up- and downregulated proteins using DAVID default annotation categories. **(H)** Scatter plot, +Dox and 24h DPI versus +Dox control. **(I)** The top 3 clusters out of 3 following functional annotation clustering of up- and downregulated proteins using DAVID default annotation categories.

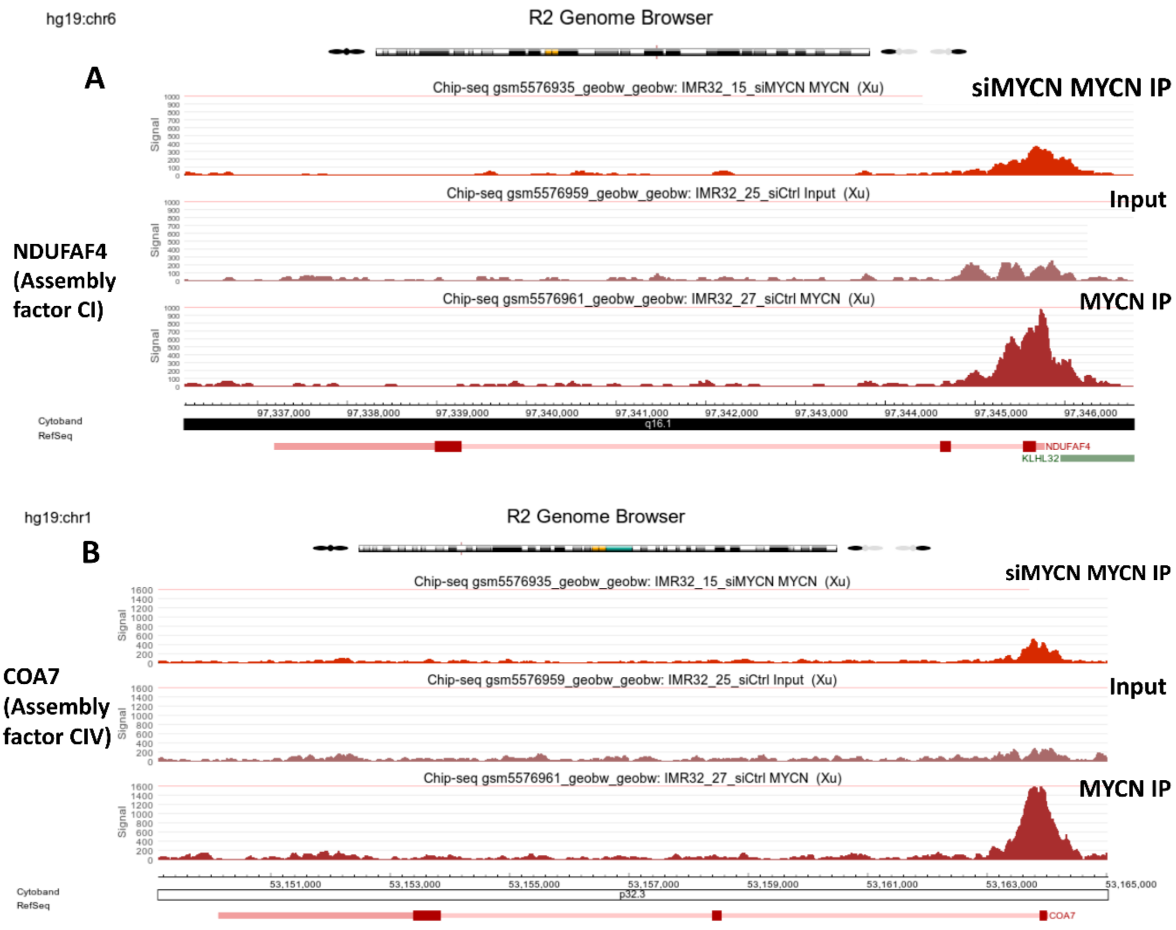


***Supplementary Figure S5. MYCN knockdown reduces promoter occupancy at NDUFAF4 and COA7 in IMR-32 NB cells.*** ChIP-seq data from a MNA NB cell line IMR-32 (*GSE183641* dataset) showing decreased occupancy at the promoter regions of **(A)** NDUFAF4 (complex I nuclear subunit gene) and **(B)** COA7 (complex IV nuclear subunit gene) (~3-fold and ~8-fold reduction, respectively) upon MYCN knockdown with siMYCN, supporting direct transcriptional regulation by MYCN. Made on R2 platform (R2: Genomics Analysis and Visualization Platform).

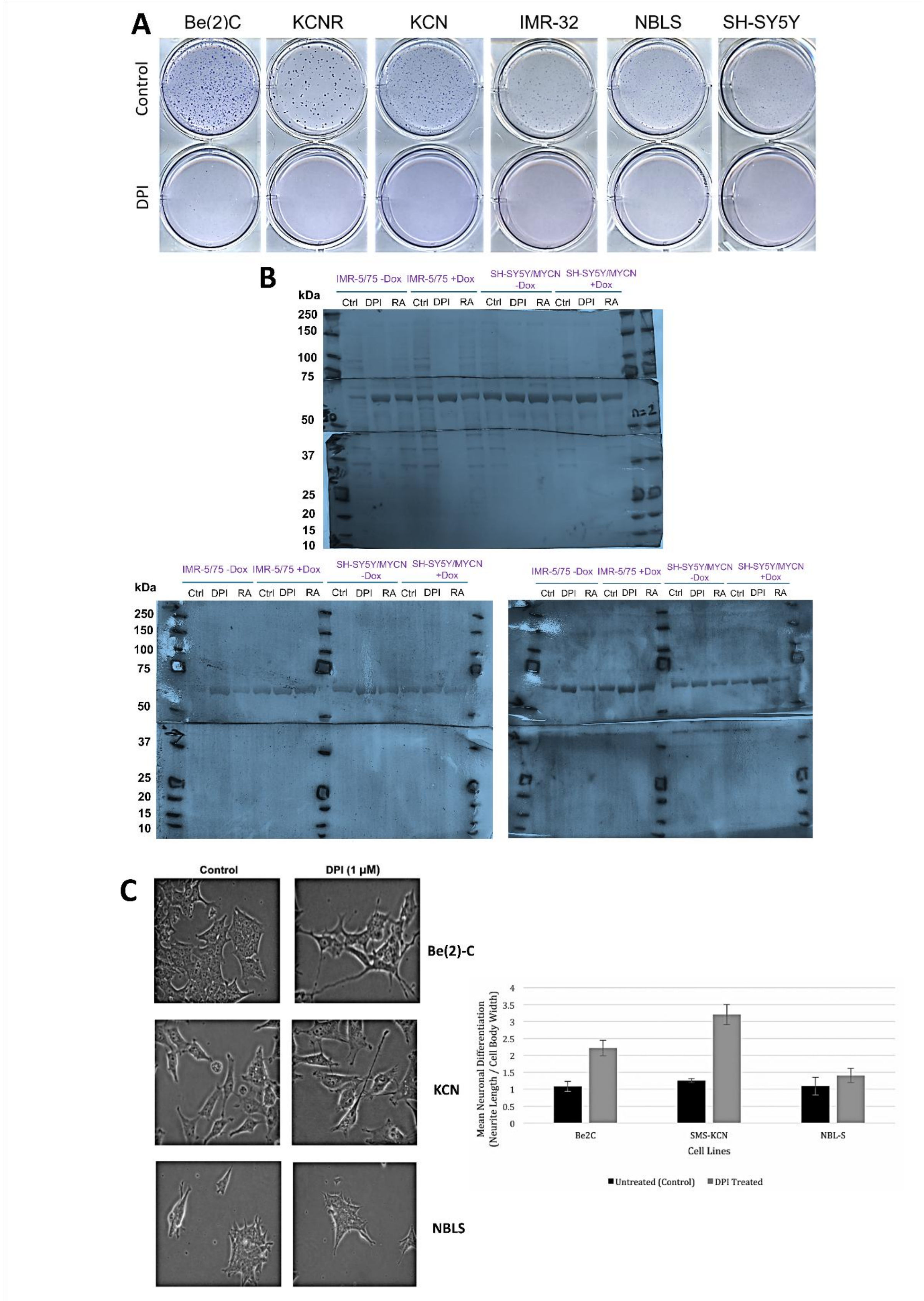


***Supplementary Figure S6. DPI decreases anchorage-independent growth and induces differentiation of NB cells.*** NB cell lines were subjected to soft agar assay. 10000 cells in 0.48% agarose were layered on a solidified 0.8% base agarose layer per well in 6-well plates.

In the presence or absence of 1µM DPI, plates were monitored for colony formation. Media +/- DPI was replenished every 4 days. After 20 days colonies were stained with crystal violet. **(B)** Coomassie blue staining of 3 membranes (3 biological replicates) showing total protein profiles from IMR-5/75 and SH-SY5Y/MYCN (-Dox/+Dox) after 5 days of either 0.5 µM DPI, 10 µM RA, or vehicle treatment. Staining confirms equal protein loading across sample**s. (C)** Phenotypic observations of Be(2)-C, SMS-KCN and NBL-S cells treated with 1 µM DPI for 72 hours. (Light microscope, X40 magnification). Longest axon of a cell / cell body width was the measure of differentiation. Measurements were calculated using ImageJ. Data represents the mean (+/- SEM) neuroblast differentiation ratio of individual cells from three technical replicates.
